# Initiation of ERAD by the bifunctional complex of Mnl1 mannosidase and protein disulfide isomerase

**DOI:** 10.1101/2024.10.17.618908

**Authors:** Dan Zhao, Xudong Wu, Tom A. Rapoport

**Author notes:** Corresponding authors: Tom Rapoport, Howard Hughes Medical Institute and Department of Cell Biology, Harvard Medical School, 240 Longwood Avenue, Boston, MA 02115, USA.,; Xudong Wu, Westlake Laboratory of Life Sciences and Biomedicine, Hangzhou, Zhejiang 310024, China.

## Abstract

Misfolded glycoproteins in the endoplasmic reticulum (ER) lumen are translocated into the cytosol and degraded by the proteasome, a conserved process called ER-associated protein degradation (ERAD). In *S. cerevisiae*, the glycan of these proteins is trimmed by the luminal mannosidase Mnl1 (Htm1) to generate a signal that triggers degradation. Curiously, Mnl1 is permanently associated with protein disulfide isomerase (Pdi1). Here, we have used cryo-electron microscopy, biochemical, and *in vivo* experiments to clarify how this complex initiates ERAD. The Mnl1-Pdi1 complex first de-mannosylates misfolded, globular proteins that are recognized through a C-terminal domain (CTD) of Mnl1; Pdi1 causes the CTD to ignore completely unfolded polypeptides. The disulfides of these globular proteins are then reduced by the Pdi1 component of the complex, generating unfolded polypeptides that can be translocated across the membrane. Mnl1 blocks the canonical oxidative function of Pdi1, but allows it to function as the elusive disulfide reductase in ERAD.

## Introduction

Newly synthesized luminal ER proteins undergo quality control to ensure that only properly folded proteins become resident in the ER or are moved on along the secretory pathway. Proteins that cannot reach their native folded state are ultimately retro-translocated into the cytosol, polyubiquitinated, and degraded by the proteasome, a conserved pathway termed luminal ER-associated protein degradation (ERAD-L) (for reviews, see ref. ^1–4^).

ERAD-L is best understood for misfolded N-glycosylated proteins in *S. cerevisiae*. The glycan is first trimmed by glucosidases and the mannosidase Mns1 to generate a Man8 species (**Fig. 1a**) ^3,5^. If the glycoprotein does not reach its native folded state, it is further processed by the mannosidase Mnl1 (also called Htm1) that generates a Man7 species containing an exposed α1,6-mannose residue (**Fig. 1a**) ^6–10^. This processing step commits the misfolded glycoprotein to ERAD-L (**Extended Data Fig. 1a**), as it is now recognized by the Hrd1 complex through interactions of the α1,6-mannose residue with the Yos9 component, and of an adjacent unstructured polypeptide segment with the Hrd3 component ^11–15^. In the next step, the polypeptide inserts as a loop into the membrane-spanning components of the Hrd1 complex; one part of the hairpin interacts with the ubiquitin ligase Hrd1, the other with the rhomboid-like protein Der1, and the tip of the loop moves through a thinned membrane region located between the two proteins ^15,16^. The hairpin likely slides through the Hrd1-Der1 interface until a suitable lysine can be polyubiquitinated by Hrd1. Finally, the protein is extracted from the membrane by the Cdc48 ATPase complex and degraded by the proteasome ^3,5^(**Extended Data Fig. 1a**).

**Figure 1.**
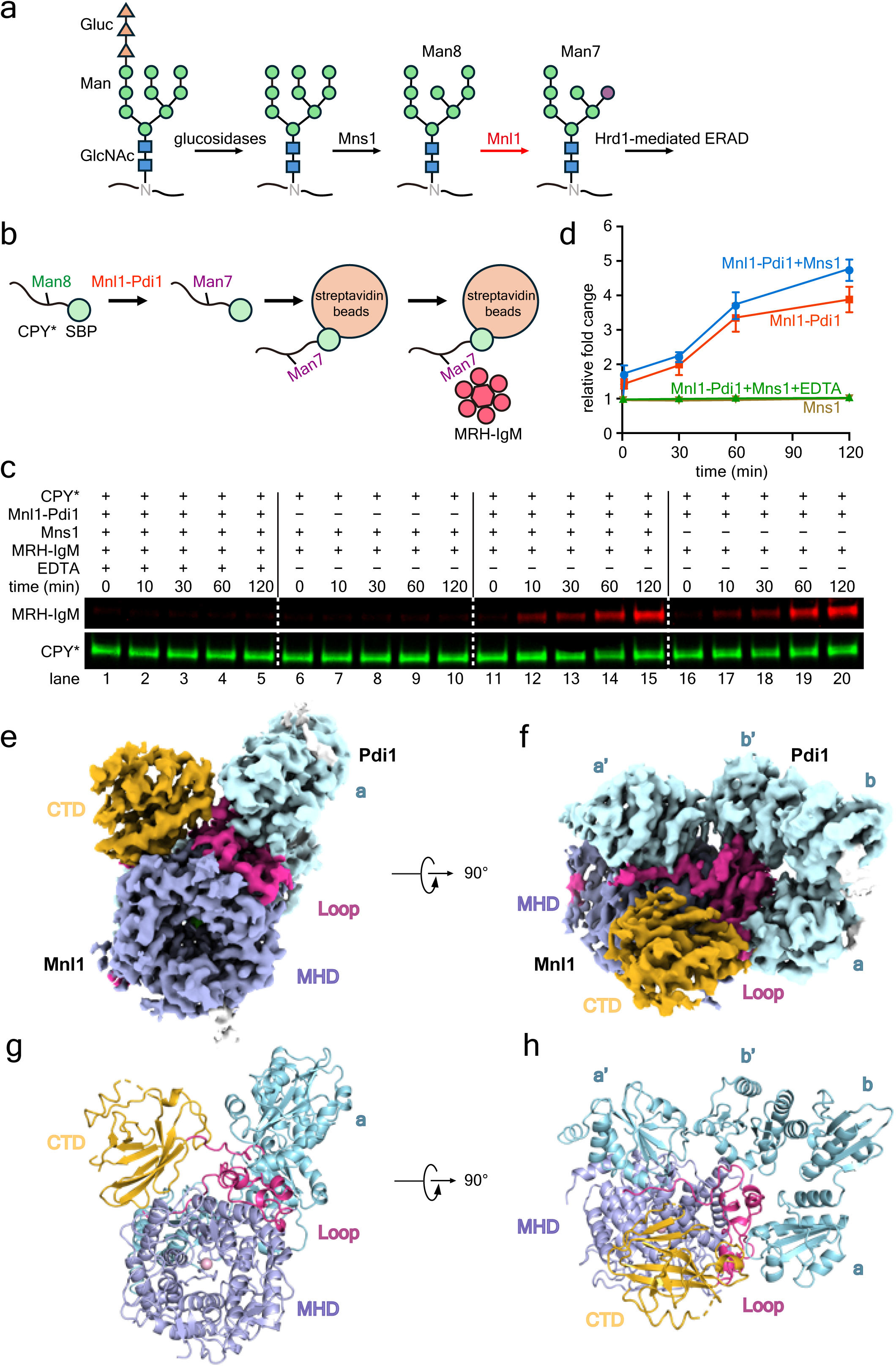
Purification and Cryo-EM structure of enzymatically active Mnl1-Pdi1 complex. **(a)** Scheme of glycan processing of a misfolded glycoprotein during ERAD-L. The conversion of Man8 to Man7 by Mnl1 commits the protein to Hrd1-mediated ERAD. The exposed α1,6-linked mannose is highlighted in purple. **(b)** Scheme showing the rational of the novel mannosidase assay. DyLight 800-labeled SBP- tagged CPY* is incubated with Mnl1-Pdi1 complex and then bound to streptavidin beads. After washing, the beads are incubated with a DyLight 680-labeled fusion of the MRH domain of OS9 and IgM (MRH-IgM). The amounts of CPY* and bound MRH-IgM are determined by SDS- PAGE and fluorescence scanning at two different wavelengths. **(c)** Mannosidase assays were performed in the presence of the indicated components. **(d)** Quantification of experiments as shown in (c) (see “Mannosidase assays” in the Methods). Shown are means and standard deviation of three experiments. **(e)** Cryo-EM density map of the Mnl1-Pdi1 complex. The mannosidase domain (MHD), the Mnl1 loop, and the C-terminal domain (CTD) of Mnl1 are shown in different colors. In this view, only Trx domain **a** of Pdi1 is visible (in cyan). **(f)** As in (e), but in a view where all Trx domains are visible. **(g)** As in (e), but with the model shown in cartoon. A Ca2+ ion is bound in the center of the MHD. **(h)** As in (f), but with a cartoon model.

The ERAD-L commitment step catalyzed by Mnl1 is not well understood. In one model, Mnl1 serves as a timer: the cleavage of the critical mannose would be slow, so that only misfolded glycoproteins that linger too long in the ER would be processed. The conversion rate of the Man8 to the Man7 glycan is indeed slow and some data show that folded proteins can be processed by Mnl1 (ref. ^17^), suggesting that they normally escape degradation by rapid vesicular export from the ER. However, many resident ER proteins have Man8 glycans.

Furthermore, folded carboxypeptidase Y (CPY) retained in the ER is processed less efficiently by Mnl1 than a misfolded variant (CPY*), and *in vitro* experiments show that Mnl1 has a preference for unfolded glycoproteins ^18^. How Mnl1 would recognize the unfolded state of a glycoprotein and distinguish terminally misfolded glycoproteins from abundant folding intermediates in the ER lumen is unclear.

Curiously, Mnl1 forms a permanent complex with protein disulfide isomerase (Pdi1) ^8,17–20^, a redox enzyme found in all eukaryotic cells. Pdi1 is more abundant than Mnl1, so only a small fraction (<10%) is found in the complex. Pdi1 is normally responsible for oxidizing cysteines to disulfides in newly synthesized proteins ^21,22^. In this process, it transfers an intramolecular disulfide bond to the substrate and becomes reduced; it is then re-oxidized by the oxygen- utilizing enzyme Ero1 ^23–27^. Pdi1 can also isomerize disulfides by transiently reducing them, and it can serve as a chaperone independent of its redox activities ^28^.

It is unclear why Mnl1 and Pdi1 are associated with one another. One possibility is that Pdi1 helps with the selection of substrates or facilitates the mannosidase reaction catalyzed by Mnl1. Another, not mutually exclusive, possibility is that Mnl1 modifies the redox activities of Pdi1. The most interesting possibility is that Pdi1 in the Mnl1-Pdi1 complex reduces disulfide bonds in misfolded proteins to facilitate their retro-translocation across the ER membrane. Such disulfide reduction has been postulated for a long time ^29–32^, and it is indeed difficult to envision that polypeptides move through the retrotranslocon as globular structures containing disulfides. However, the identity of the reductase has been elusive. In mammals, disulfides can be reduced by the PDI homolog ERdj5 ^33,34^, but this enzyme does not exist in yeast. Although Pdi1 and its mammalian homolog PDI play a role in ERAD ^31,35,36,37^, their exact role has yet to be established.

In this paper, we clarify how the Mnl1-Pdi1 complex initiates ERAD in *S. cerevisiae*. We show that the complex first trims the glycan of misfolded, globular proteins and then reduces the disulfides, generating unfolded polypeptides that can be retro-translocated across the membrane. Our results indicate that Pdi1 in the Mnl1-Pdi1 is the elusive reductase involved in ERAD.

## Results

### Architecture of the Mnl1-Pdi1 complex

To better understand the function of the Mnl1-Pdi1 complex, we first determined a cryo-EM structure. The Mnl1-Pdi1 complex was purified from *S. cerevisiae* cells that overexpress a FLAG-tagged version of Mnl1 (Mnl1-FLAG) together with Pdi1. The complex was released from the lumen of a membrane fraction by detergent treatment, enriched with beads containing FLAG antibodies, and further purified as a 1:1 complex by size-exclusion chromatography (Extended Data Fig. 1b,c).

The purified Mnl1-Pdi1 complex was enzymatically active, as it could generate the α1,6- mannose residue in the glycan of the misfolded glycoprotein CPY*, an established ERAD-L substrate ^38,39^. We measured the mannosidase activity with a novel assay that circumvents the previous cumbersome analysis by HPLC and mass spectrometry. The purified Mnl1-Pdi1 complex was first incubated with a 10-fold molar excess of DyLight 800-labeled streptavidin-binding peptide (SBP)-tagged CPY* (CPY*-SBP). After retrieval of CPY*-SBP with streptavidin beads, the generation of the α1,6-mannose residue was determined by the binding of DyLight 680-labeled mannose 6-phosphate receptor homology (MRH) domain of OS9 (the mammalian homolog of Yos9), fused to oligomeric immunoglobulin M (MRH-IgM) (see scheme in **Fig. 1b**). The bound material was eluted with biotin and analyzed by SDS-PAGE followed by fluorescence scanning at two different wavelengths. The Man7 glycan was generated whether or not the upstream enzyme Mns1 was present (**Fig. 1c**, lanes 11-15 versus 16-20, and **Fig. 1d**), suggesting that purified CPY*-SBP already contains Man8 glycans and that the Mnl1-catalyzed reaction is rate-limiting. As expected, Mns1 alone did not generate the Man7 species (lanes 6- 10), and the reaction was inhibited by EDTA, which chelates the essential Ca^2+^-ions ^18,20^(lanes 1- 5), or by mutation of an active site residue in Mnl1 (D279N) (**Extended Data Fig. 1d**).

A cryo-EM structure of the Mnl1-Pdi1 complex was obtained from a particle class after 3D classification and refinement and had an overall resolution of 3.0 Å (**Fig. 1e-h** and **Extended Data Fig. 2**). The density map of this class allowed model building for most parts of the proteins (see examples in **Extended Data Fig. 3a-d**), with the exception of some Mnl1 loops. The density of a C-terminal domain (CTD) was also weaker than that for other parts of the protein (**Extended Data Fig. 3d**). Another major 3D class resulted in a similar, lower quality, density map at 3.2Å overall resolution, but it lacked the CTD (**Extended Data Fig. 2** and **Extended Data Fig. 3e,f**). Thus, the CTD of Mnl1 seems to be rather flexible.

Mnl1 consists of a canonical mannosidase domain (MHD; amino acids 29-514), a loop interacting with Pdi1 (residues 515-650), and the CTD (residues 651-796) (**Fig. 1e-h**). The MHD contains α-helices that form a barrel with a central pore (**Fig. 1e,g**). This domain is superimposable with that of Mns1 (**Extended Data Fig. 4a**), which lacks all the other domains of Mnl1 ^40^. As shown for Mns1 ^40^, the central pore contains the active site residues and the essential Ca^2+^ ion (**Fig. 1g** and **Extended Data Fig. 3c**); it accommodates the glycan of the glycoprotein substrate during the mannosidase reaction. The CTD interacts with one side of the mannosidase barrel using a relatively small interface (**Fig. 1g**). Pdi1 contains four thioredoxin-like (Trx) domains, referred to as **a**, **b**, **b’**, and **a’** domains, with **a** and **a’** containing redox-active CGHC motifs. The Mnl1 loop interacts with three of the four Trx domains of Pdi1, i.e. domains **a**, **b’**, and **a’** (**Fig. 1h**). The total interaction surface is fairly large (2,480Å^2^). The four Trx domains of Pdi1 form a U shape that is most similar to the structure of the reduced form of human PDI ^41^ (**Extended Data Fig. 4b**).

### Disulfide bonds between Mnl1 and Pdi1

The Mnl1 loop contains two cysteines (Cys579 and Cys 644) that are in disulfide-bonding distance to the first cysteines of the redox-active CGHC motifs of the **a’** and **a** Trx domains of Pdi1 (**Fig. 2a,b**). The density map supports the formation of disulfide bonds (**Extended Data Fig. 3a,b**), even though a covalent adduct could not be detected in non-reducing SDS-PAGE (**Extended Data Fig. 1c**). In intact yeast cells, a sizable fraction of FLAG-tagged Mnl1, expressed at endogenous levels, formed disulfide-linked complexes that contain Pdi1 (**Extended Data Fig. 1e**), consistent with data in the literature ^19^. In addition, the adducts seem to contain a heterogeneous mixture of substrate molecules that are likely disulfide-linked to the Pdi1 component. Taken together, these results suggest that the **a** and **a’** domains of Pdi1 form reversible disulfide bonds with the Mnl1 loop. However, the disulfide bonds are not required for the interaction between the two proteins, because FLAG-tagged Mnl1 co-precipitated Pdi1 even when all cysteines were reduced with dithiothreitol (DTT) (**Supplementary Fig. 1a**).

**Figure 1.**
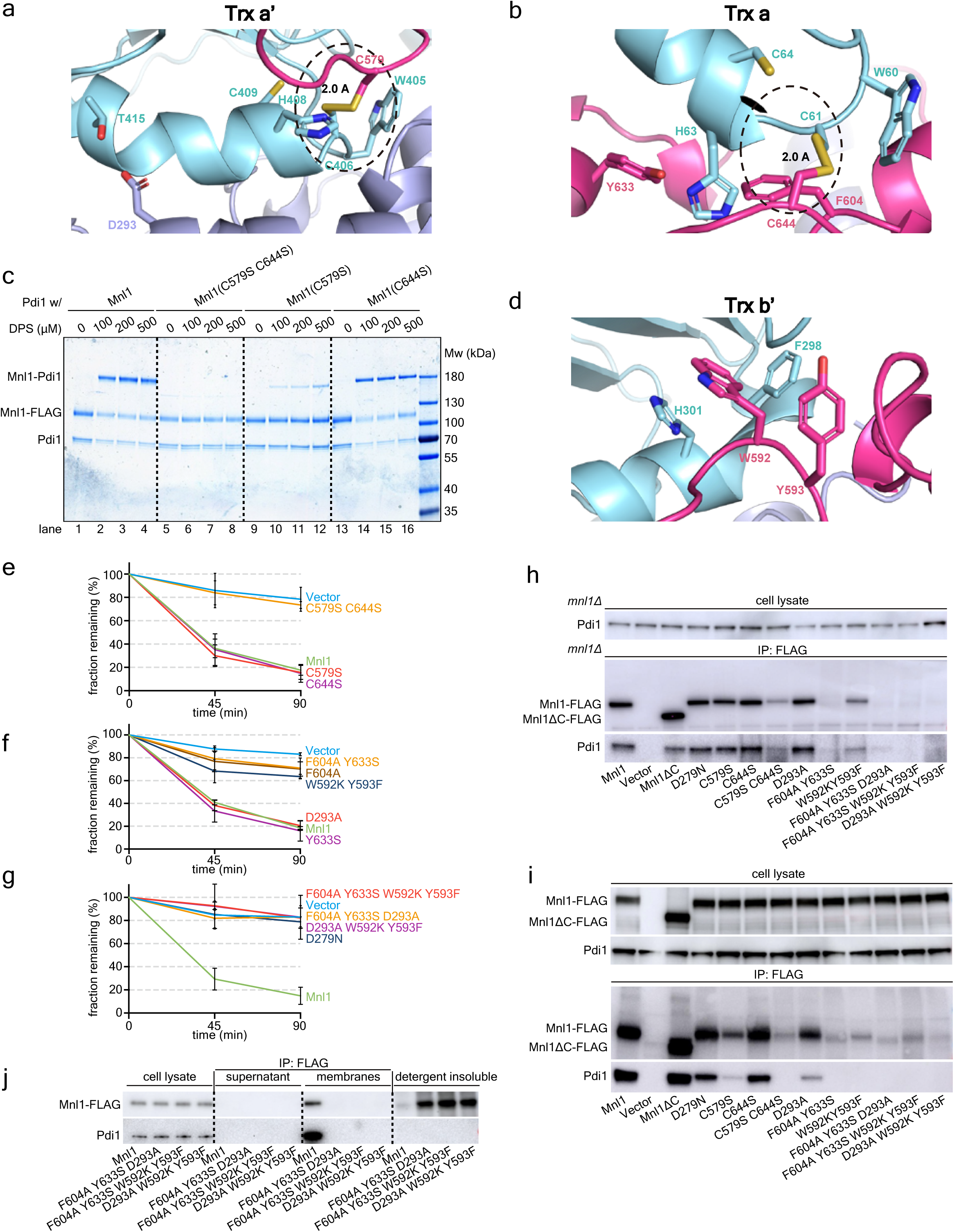
Interactions between Mnl1 and Pdi1. **(a)** Cysteine C579 in the Mnl1 loop forms a disulfide bond with the first cysteine (C406) of the CGHC motif of the Trx **a’** domain of Pdi1 (encircled with a broken line). **(b)** As in (a), but for cysteine C644 of the Mnl1 loop and the first cysteine (C61) of the CGHC motif of the Trx **a** domain. **(c)** Purified complexes of Pdi1 with wild-type Mnl1 or the indicated cysteine mutants were incubated with different concentrations of 2,2ʹ- dipyridyldisulfide (DPS) to induce disulfide bridge formation between Pdi1 and Mnl1. The samples were subjected to non-reducing SDS- PAGE and staining with Coomassie blue. **(d)** Residues of the Mnl1 loop inserted into the hydrophobic pocket of the Trx **b’** domain. **(e)** ERAD of CPY*-HA was determined by cycloheximide (CHX) chase experiments in cells lacking Mnl1 (*mnl1*Δ). The cells were transformed with either an empty vector or expressed wild-type Mnl1 or the indicated cysteine mutants. The samples were analyzed by SDS-PAGE and immunoblotting for HA. The intensities of the CPY*-HA bands were quantified. Shown are the fractions of CPY*-HA remaining at different time points (means and standard deviation of three experiments). **(f)** As in (e), but with other residues mutated at the interface between Mnl1 and Pdi1. **(g)** As in (f), but with additional interface mutants. **(h)** The indicated FLAG-tagged Mnl1 mutants were expressed from the endogenous locus. A membrane fraction was solubilized in Nonidet P-40 and the extract subjected to immunoprecipitation (IP) with FLAG antibodies. The samples were analyzed by SDS-PAGE and immunoblotting for FLAG and Pdi1. A sample of the cell lysate was analyzed directly for Pdi1. **(i)** As in (h), but with overexpressed Mnl1-FLAG constructs. A sample of the cell lysate was analyzed by immunoblotting for FLAG and Pdi1. Note that all Mnl1 versions were expressed at about equal levels. **(j)** As in (i), but cell lysates were separated into supernatant and membrane fractions. Membrane fractions were incubated with Nonidet P-40 and centrifuged again. The supernatants and detergent extracts were subjected to immunoprecipitation with FLAG antibodies. All samples were analyzed by SDS-PAGE and immunoblotting for FLAG and Pdi1.

Disulfide bond formation between Mnl1 and Pdi1 is supported by experiments in which we added increasing concentrations of 2,2’-dipyridyldisulfide (DPS) and subjected the samples to non-reducing SDS-PAGE (**Fig. 2c**). Adduct formation was observed with wild-type Mnl1 (lanes 1- 4) or mutants in which one of the two cysteines (Cys579 and Cys644) of the Mnl1 loop were mutated (lanes 9-12 and 13-16), although it was less efficient with the Cys579 mutant. As expected, when both cysteines were mutated together, no adduct was formed (lanes 5-8). Superimposing the active sites of the **a** and **a’** domains shows that they bind Mnl1 segments in a similar manner (**Extended Data Fig. 4c**).

Mutation of each of the two cysteines in the Mnl1 loop individually had little effect on ERAD-L of CPY*-HA, but mutation of both together abolished degradation (**Fig. 2e** and **Supplementary Fig. 1b**). Thus, one of the two possible disulfide bonds with Pdi1’s active sites is required for Mnl1 function. However, the redox state of Pdi1 does not affect the mannosidase activity of Mnl1, as shown by adding different ratios of oxidized and reduced glutathione (**Extended Data Fig. 1f**), or by adding DPS to force disulfide bridge formation (**Extended Data Fig. 1g**).

Our Mnl1-Pdi1 structure shares several features with those of other ER proteins that form stable complexes with PDI or its homologs (**Extended Data Fig. 4d-j**). These include the prolyl 4- hydroxylases (C-P4H) involved in collagen synthesis ^42^, the microsomal triglyceride transfer protein (MTP) involved in lipoprotein assembly ^43^, and the tapasin protein involved in peptide loading onto the MHC class I molecule, which forms a complex with the PDI-homolog ERp57 ^44^. The four Trx domains of the redox partners always adopt a U shape in which the **a** and **a’** domains interact with the client protein, and a cysteine in the client is always close to the first cysteine in one of the active site CXXC motifs. In the case of collagen 4-prolylhydroxylase, two cysteines are positioned next to the CXXC motifs, as in the Mnl1-Pdi1 complex (**Extended Data Fig. 4i,j**).

### Pdi1 keeps Mnl1 soluble in the ER lumen

A hydrophobic pocket of the **b’** Trx domain accommodates Trp592 and Tyr593 of the Mnl1 loop (**Fig. 2d**), similarly to how this domain in other structures interacts with hydrophobic amino acids ^42,43^. Mutations in the loop designed to disrupt this interaction reduced Mnl1’s function in ERAD-L (**Fig. 2f** and **Supplementary Fig. 1c**). A role for the **b’** domain is also supported by *in vivo* data with a pdi1 allele, pdi1-1, that carries a Leu313Pro mutation near the hydrophobic pocket^20^. The Mnl1 loop also binds to the **a** domain (**Fig. 2b**), and this interaction is important for ERAD, as shown by mutating Phe604 (**Fig. 2f** and **Supplementary Fig. 1c**). A mutation designed to disrupt the interaction with the **a’** domain did not affect ERAD (D293A; **Fig. 2a,f** and **Supplementary Fig. 1c**), but combining the mutations that disrupt the **a** or **b’** interface with other mutations or mutating an active site residue in the mannosidase domain (D279N) resulted in ERAD-defective Mnl1 variants (**Fig. 2g** and **Supplementary Fig. 1d**).

Mnl1 mutants that cannot interact with Pdi1 form insoluble aggregates in the ER. When FLAG-tagged Mnl1 was expressed from the endogenous promoter in *S. cerevisiae* cells, all mutants defective in ERAD showed reduced levels in the detergent-solubilized membrane fraction after immunoprecipitation (**Fig. 2h**). Similar results were obtained when Mnl1-FLAG was overexpressed, so that it could be detected in crude cell lysates by immunoblotting (**Fig. 2i**); all mutants were expressed at about the same level as the wild-type protein, but little could be solubilized with detergent (**Fig. 2j**). Small amounts of the mutant complexes with disturbed interfaces (W592K, Y593F and C579S, C644S mutants) could be purified (**Extended Data Fig. 5a**) and had significant mannosidase activity (**Extended Data Fig. 1d**), demonstrating that Pdi1 is primarily required to keep Mnl1 soluble in the ER lumen. Because Mnl1 lacks an ER retention signal, Pdi1 may also localize the mannosidase to the ER.

### The CTD in the Mnl1-Pdi1 complex recognizes misfolded, globular proteins

Next we asked how substrates are recognized by the Mnl1-Pdi1 complex. We suspected that misfolded ERAD substrates are recruited by the CTD, as a mutant lacking the CTD (Mnl1ΔC) had only low mannosidase activity with CPY* as the substrate (**Fig. 3a**, lanes 7-9 versus 1-3, and **Fig. 3b**) and was inactive in ERAD (**Fig. 3c** and **Supplementary Fig. 1e**). The mutant retained the interaction with Pdi1, allowing the purification of a stable complex (**Extended Data Fig. 5b**), and formed a disulfide-linked adduct after addition of DPS (**Extended Data Fig. 5c**).

**Figure 3.**
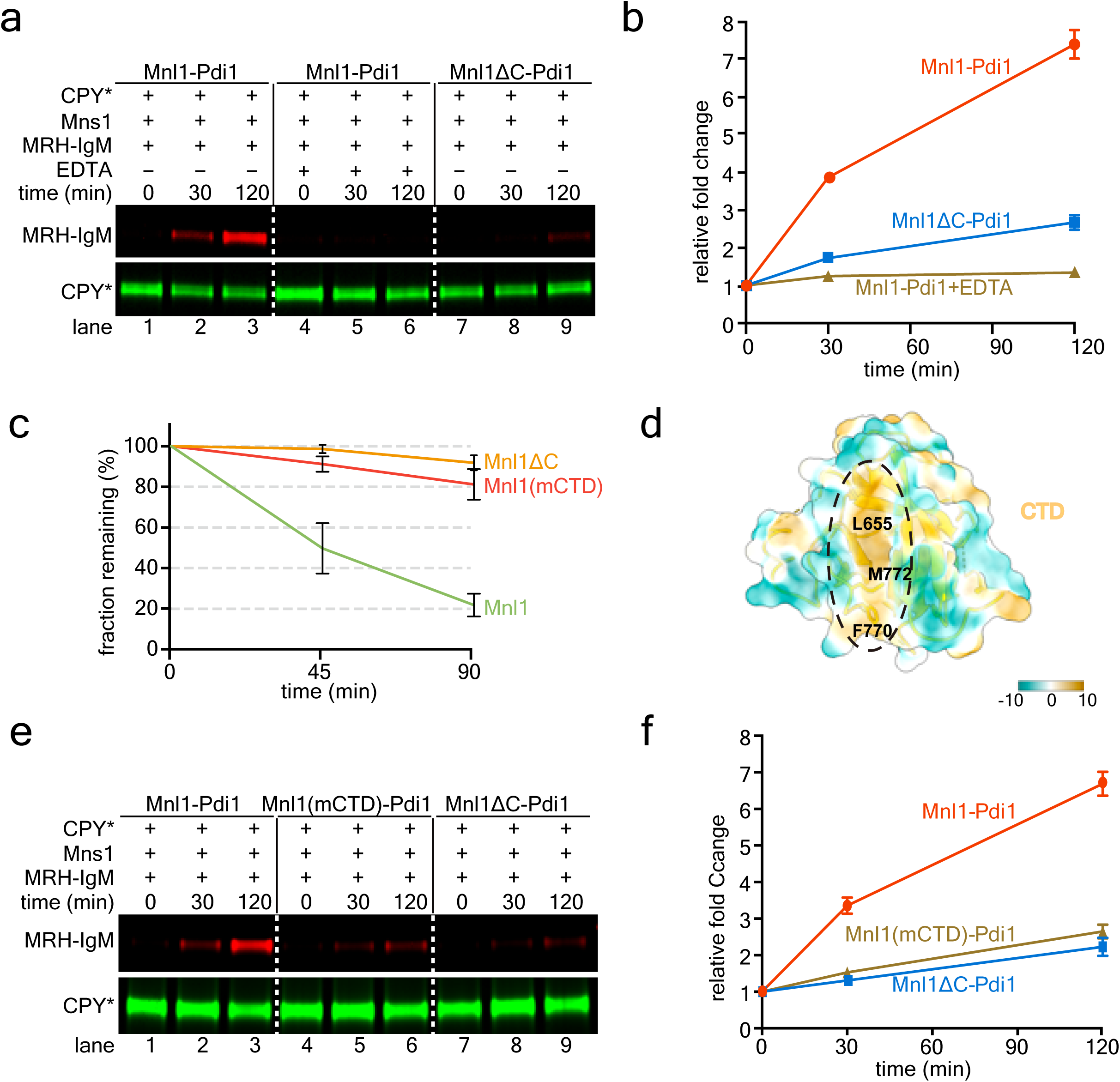
The role of Mnl1’s C-terminal domain (CTD) in ERAD. **(a)** Mannosidase assays were performed with purified wild-type or mutant Mnl1-Pdi1 complex. **(b)** Quantification of experiments as in (a) (see “Mannosidase assays” in the Methods). Shown are means and standard deviation of three experiments. **(c)** ERAD of CPY*-HA was determined by cycloheximide (CHX) chase experiments in cells lacking Mnl1 (*mnl1*Δ). The cells were transformed with wild-type Mnl1 or the indicated mutants. The samples were analyzed by SDS-PAGE and immunoblotting for HA. The intensities of the CPY*- HA bands were quantified. Shown are the fractions of CPY*-HA remaining at different time points (means and standard deviation of three experiments). **(d)** Hydrophobic groove of the CTD (hydrophobicity scale on the right). Shown is a semi- transparent space-filling model with a cartoon model. The three hydrophobic residues mutated in the mCTD mutant are labeled. **(e)** Mannosidase assays were performed with the indicated purified wild-type or mutant Mnl1- Pdi1 complexes. **(f)** Quantification of experiments as in (e). Shown are means and standard deviation of three experiments.

The CTD has a pronounced hydrophobic groove located between two β-sheets (**Fig. 3d** and **Extended Data Fig. 3d**). We generated a mutant (mCTD), in which three hydrophobic residues in the groove were changed to hydrophilic amino acids (L655N, F770Y, M772K). The mutant was defective in ERAD (**Fig. 3c** and **Supplementary Fig. 1e**), and the purified complex with Pdi1 (see **Extended Data Fig. 5d**) had reduced mannosidase activity (**Fig. 3e**, lanes 4-6 versus 1-3, and **Fig. 3f**). These data suggest that the hydrophobic groove of the CTD binds exposed hydrophobic segments in misfolded ERAD substrates.

Next we tested the mannosidase activity of the Mnl1-Pdi1 complex with RNase B (RB) versions of different folding status. RB differs from the well-characterized RNase A (RA) by the presence of an N-glycan ^18^ (**Fig. 4a**). A non-native conformation of RB was obtained by the removal of the N-terminal S-peptide (RBΔS) ^45^. A completely unfolded state (RBun) was generated by treating RB with guanidinium hydrochloride and dithiothreitol (DTT), followed by the removal of the denaturants on a desalting column ^18^. Finally, a more compact misfolded state was generated from RBun by oxidizing the cysteines with diamide to form random disulfides (scrambled RB; RBsc) ^18^ (**Fig. 4a**). The misfolded states of RBΔS and RBsc probably resemble the partially folded conformations of many ERAD substrates. Folded RB was not a mannosidase substrate for Mnl1-Pdi1 (**Fig. 4b**, lanes 1-4), demonstrating that the complex recognizes the misfolded state of a protein. Furthermore, RBsc and particularly RBΔS, were much better mannosidase substrates than RBun (**Fig. 4b**, lanes 13-16 and 5-8 versus lanes 9- 12). These results indicate that Mnl1 preferentially processes the glycan of partially folded, globular proteins, as reported previously ^18^.

**Figure 4.**
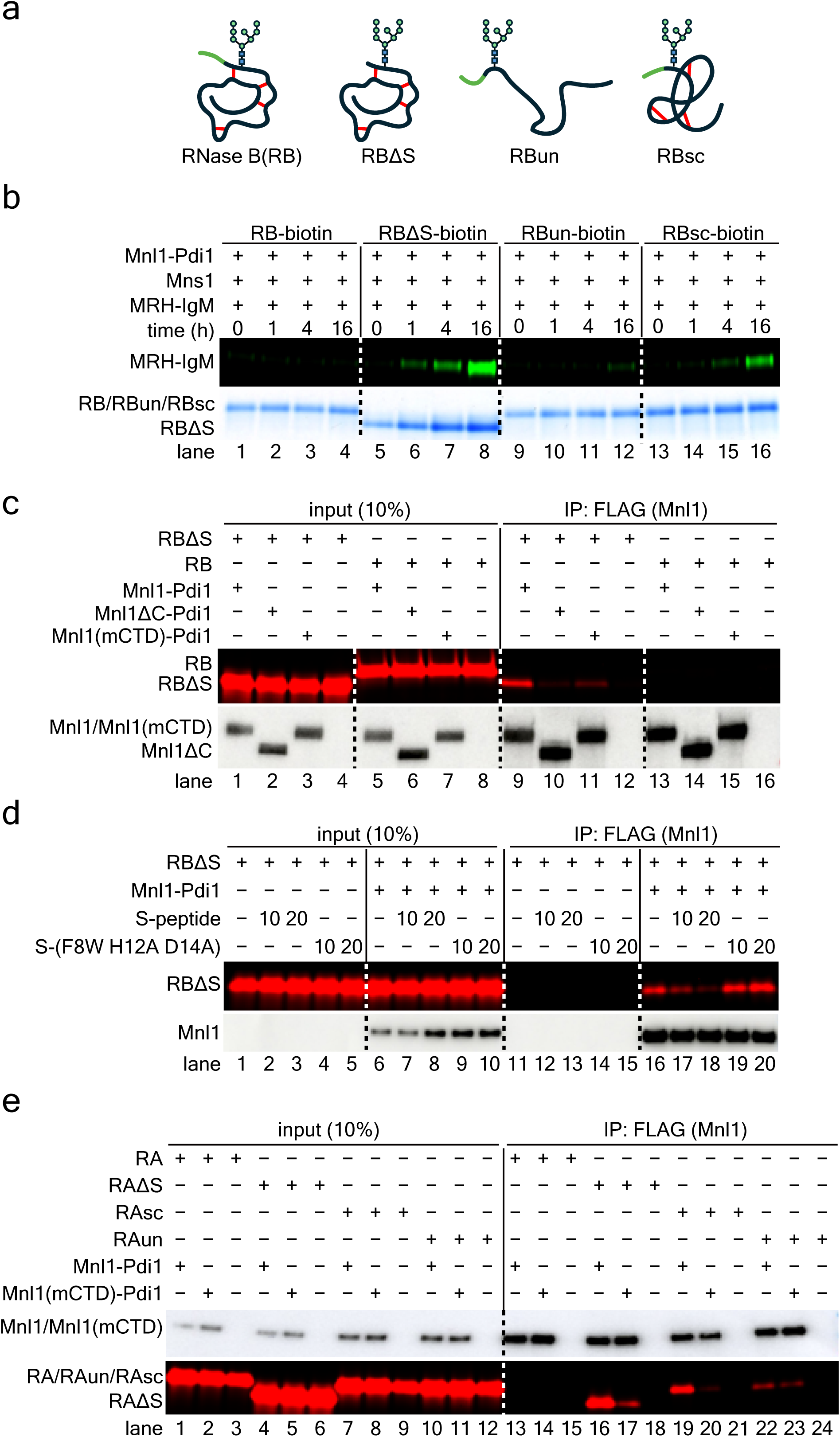
The Mnl1-Pdi1 complex interacts with misfolded polypeptides. **(a)** Schematic of different versions of misfolded RNase B. RB, folded RNase B; RBΔS, RB lacking the S-peptide; RBun, completely unfolded RNase B; RBsc, RB with scrambled disulfides. The S-peptide is colored in green, and disulfide bonds are shown as red lines. **(b)** Mannosidase reactions were performed with Mnl1-Pdi1 complex and the indicated substrates labeled with biotin. After the mannosidase reaction, the substrates were retrieved with streptavidin beads and the binding of DyLight 800-labled MRH-IgM was determined. The amount of substrate in the assays was monitored by Coomassie blue staining. **(c)** Wild-type or mutant Mnl1-Pdi1 complex was incubated with DyLight 680-labeled RBΔS or RB. The complexes were retrieved with beads containing FLAG antibodies and bound substrate analyzed by SDS-PAGE and fluorescence scanning. The amounts of Mnl1-FLAG in the assays were determined by immunoblotting for FLAG. A fraction of the input was analyzed directly. **(d)** As in (c), but with RBΔS and addition of a synthetic S-peptide at 10 or 20-fold molar excess. A control was performed with a mutant S-peptide carrying three mutations (F8W H12A D14A) that prevent binding to RBΔS. **(e)** As in (c), but with folded and misfolded RNase A (RA) versions (which lack a glycan).

Pull-down experiments showed that these proteins bind to the CTD: full-length Mnl1-Pdi1 complex bound RBΔS (**Fig. 4c**, lane 9), but not RB (lane 13), whereas the complex lacking the CTD or carrying mutations in the hydrophobic pocket of CTD (mCTD) bound neither protein strongly (lanes 10 and 14, 11 and 15, respectively). The addition of increasing concentrations of a synthetic S-peptide to RBΔS restored the folded state of RB, as previously shown for RNase A ^46,47^, and consequently abolished the binding of RBΔS to Mnl1-Pdi1 (**Fig. 4d**, lanes 17, 18). In contrast, a mutant version of the S-peptide that does not interact with RBΔS ^48,49^ did not affect the binding (lanes 19, 20). These data confirm that Mnl1 recognizes the non-native state of RBΔS through its CTD. We also used folded and misfolded versions of non-glycosylated RNase A (RA, RAΔS, RAsc, and RAun). These proteins behaved like the corresponding RB variants in pull-down experiments (**Fig. 4e**, and see below), indicating that the Mnl1-Pdi1 complex recognizes the misfolded state of globular proteins, rather than the glycan.

### Pdi1 modifies the substrate specificity of the CTD

Surprisingly, we found that the isolated CTD, purified as a fusion with maltose binding protein (MBP) from mammalian tissue culture cells (MBP-CTD) (**Extended Data Fig. 5e**), preferentially bound completely unfolded polypeptides. In the absence of MBP-CTD, firefly luciferase (Luc) or citrate synthase (CiS), two established chaperone substrates ^50,51^, formed large aggregates when the incubation temperature was increased, as detected by dynamic light scattering (**Fig. 5a,b**). Aggregation was gradually prevented by increasing concentrations of MBP-CTD (**Fig. 5a,b**). In contrast, purified MBP-mCTD (**Extended Data Fig. 5e**), containing the three mutations in the hydrophobic groove, or the fusion partner MBP alone, had no effect on aggregation (**Fig. 5c,d** and **Extended Data Fig. 5f,g**). Pull-down experiments confirmed that MBP-CTD does not interact with fluorescently labeled, folded RB (**Fig. 5e**, lane 14), and binds RBun stronger than RBΔS or RBsc (lane 15 versus lanes 13 and 16), in contrast to the substrate preference of the CTD in the Mnl1-Pdi1 complex (**Fig. 4**). Thus, Pdi1 in the Mnl1-Pdi1 complex seems to cause the CTD to ignore unfolded polypeptides.

**Figure 5.**
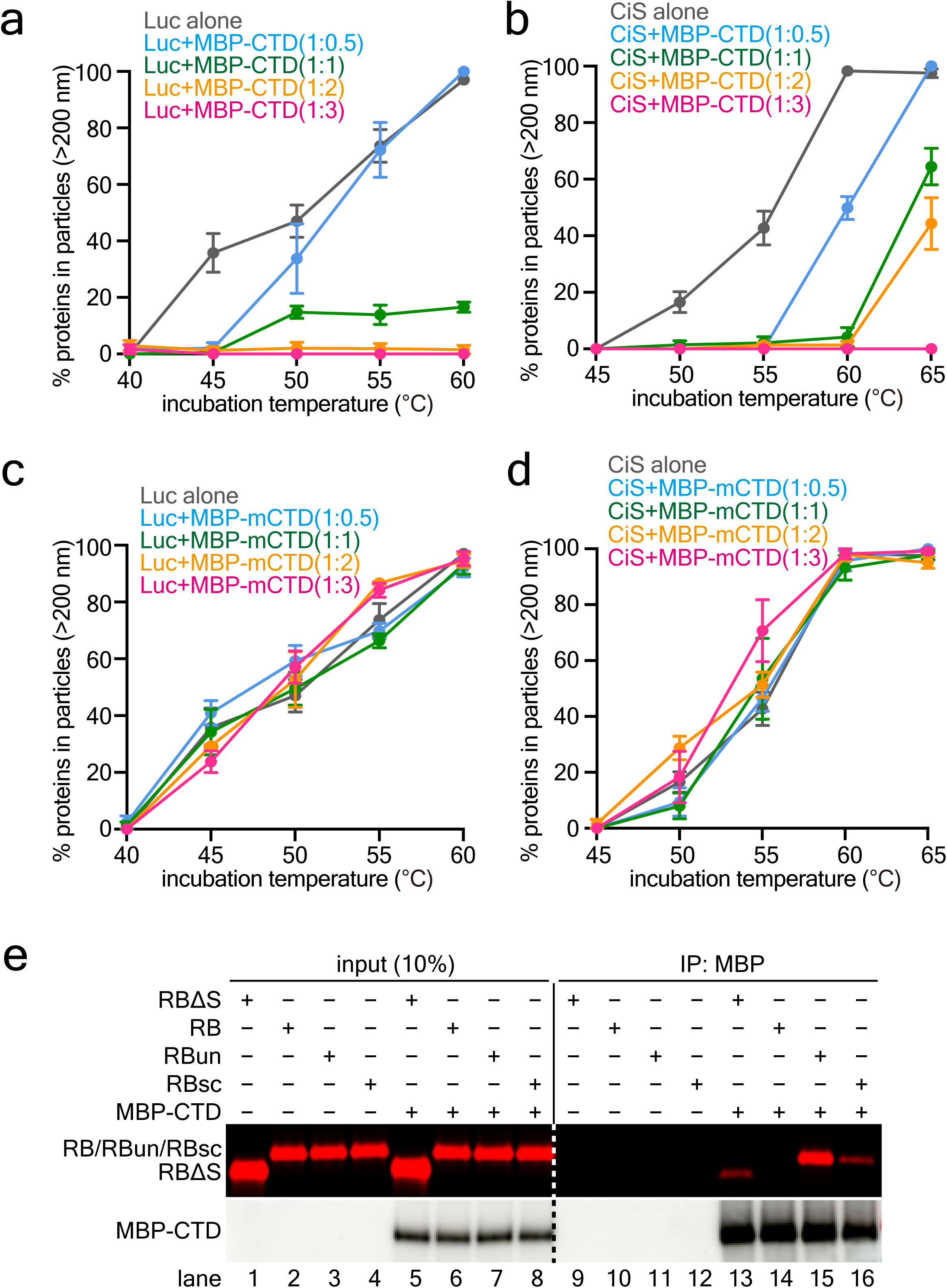
The isolated CTD of Mnl1 interacts with unfolded polypeptides. **(a)** A fusion of MBP and CTD (MBP-CTD) was incubated with luciferase (Luc) at different molar ratios for 20 min at different temperatures. A control was performed with Luc alone. The samples were analyzed by dynamic light scattering and the percentage of Luc in particles larger than 200nm (aggregates) was determined. Shown are means and standard deviations of three experiments. **(b)** As in (a), but with citrate synthase (CiS). **(c)** As in (a), but incubation of Luc with MBP-mCTD, in which the hydrophobic groove is mutated. **(d)** As in (c), but with CiS. **(e)** RB, RBΔS, RBun, or RBsc were labeled with the fluorescent dye DyLight 680 and incubated with or without MBP-CTD. Material bound to MBP-CTD was retrieved with resin containing MBP antibodies and analyzed by SDS-PAGE followed by fluorescent scanning. The samples were also analyzed by blotting for MBP. A fraction of the input material was analyzed directly.

### Pdi1 in the Mnl1-Pdi1 complex recognizes unfolded polypeptides

Pdi1 probably does not modify the behavior of the CTD directly, as they do not interact in our Mnl1-Pdi1 structure. However, it seemed possible that Pdi1 prevents the binding of unfolded polypeptides to the CTD by competing for them. Consistent with this model, the addition of RBun caused the dissociation of the Mnl1-Pdi1 complex, abolishing the formation of disulfide bonds between the two cysteines in the Mnl1 loop and the active sites of the Trx **a** and **a’** domains (**Fig. 6a**, lanes 5-8 versus 1-4, and **Extended Data Fig. 5h**). An approximately 2-fold molar was sufficient for half-maximal inhibition (**Extended Data Fig. 5h**). In contrast, the globular, misfolded substrates RBΔS and RBsc were without effect (**Fig. 6b,c**). RBun bound to Pdi1, rather than Mnl1, as it interfered with disulfide bridge formation even when the CTD was absent or mutated in its hydrophobic residues (**Extended Data Fig. 5h**). RBun could also block disulfide bond formation between Mnl1 and Pdi1 when one of the two cysteines in the Mnl1 loop was mutated (**Fig. 6a** and **Extended Data Fig. 5i**).

**Figure 6.**
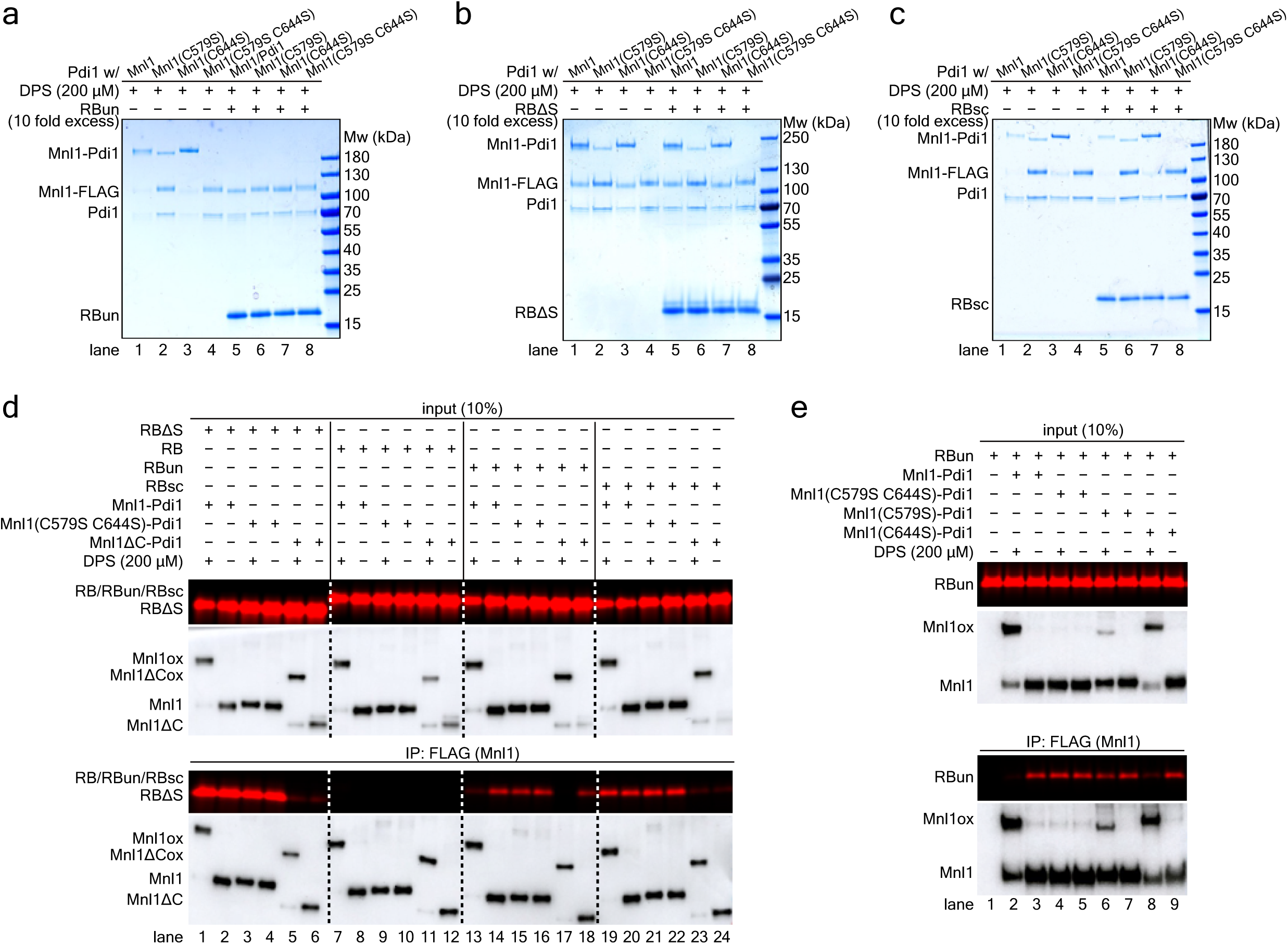
Mnl1 and Pdi1 have distinct interactions with misfolded polypeptides. **(a)** Wild-type or mutant Mnl1-Pdi1 complexes were incubated with a 10-fold molar excess of RBun, as indicated. DPS was added to induce disulfide formation and the samples were analyzed by non-reducing SDS-PAGE and Coomassie blue staining. **(b)** As in (a), but with RBΔS. **(c)** As in (a), but with RBsc. **(d)** Folded RNase B (RB) or the indicated misfolded variants were fluorescently labeled and incubated with wild-type or mutant Mnl1-Pdi1 complexes containing FLAG-tagged Mnl1 versions. Where indicated, the complexes were pretreated with DPS to induce disulfide bridges between the Mnl1 loop cysteines and the active site cysteines of Pdi1. After immunoprecipitation with FLAG antibodies, bound substrate was analyzed by SDS-PAGE and fluorescence scanning. The samples were also analyzed by immunoblotting for FLAG. A fraction of the input material was analyzed directly. **(e)** As in (d), but with RBun and the indicated cysteine mutants in the Mnl1 loop.

To further confirm that the globular, misfolded proteins RBΔS and RBsc bind to the CTD of Mnl1, whereas the fully unfolded protein RBun binds to Pdi1, we performed pull-down experiments of the Mnl1-Pdi1 complex with FLAG-tagged versions of Mnl1 (**Fig. 6d**). Indeed, the binding of RBΔS or RBsc was abolished when the CTD was deleted (lanes 2 and 20 versus 6 and 24). Pre-treatment of the Mnl1-Pdi1 complex with DPS to generate disulfides that lock the active Trx domains of Pdi1 to the Mnl1 loop had no effect (lanes 1 and 19). By contrast, the binding of RBun was only slightly affected by the deletion of the CTD (lanes 14 versus 18) and instead reduced or abolished when the Mnl1-Pdi1 complex was pre-treated with DPS (lanes 13 and 17). Some residual RBun binding was observed (lane 13 versus 17), suggesting that the CTD can interact with the unfolded polypeptide when the competing interaction with Pdi1 is abolished. As expected, DPS did not prevent the binding of RBun when both cysteines in the Mnl1 loop were mutated (lanes 15 and 16). However, RBun binding was abolished when Cys644 was mutated (C644S), consistent with this mutant still allowing efficient disulfide bond formation between Mnl1 and Pdi1 (**Fig. 6e**, lane 8 versus 2). On the other hand, only a slight effect was observed with the other single-cysteine mutant (C579S) that did not allow efficient adduct formation (lane 6 versus 2). These data support the idea that a partial dissociation of Pdi1 from Mnl1 is required for RBun binding to the complex. The distinct substrate specificities of the CTD and Pdi1 were confirmed with an assay in which we tested by microscopy the binding of fluorescently labeled RB, RBΔS, or RBun to beads containing Mnl1-Pdi1 complex (**Extended Data Fig. 6a-c**). RB did not bind (**Extended Data Fig. 6a,c**) and RBΔS bound to Mnl1-Pdi1 in a CTD-dependent manner (**Extended Data Fig. 6b,c**); the interaction was not affected by DPS treatment or mutation of the two cysteines in the Mnl1 loop. In contrast, the weaker interaction of RBun was reduced when the complex was pretreated with DPS (**Extended Data Fig. 6b,c**). Taken together, our results indicate that RBun interacts with Pdi1 in the Mnl1-Pdi1 complex and causes the dissociation of the complex when in excess.

### Mnl1 modifies the redox reactions of Pdi1

Given that misfolded, globular proteins are the preferred mannosidase substrates of the Mnl1- Pdi1 complex, any disulfides in these substrates probably need to be dissolved to generate unfolded polypeptides for subsequent retro-translocation. We therefore wondered whether the Pdi1 component of the complex could serve as a disulfide reductase.

We found that Pdi1 in the complex can no longer perform its canonical oxidative function, i.e. cooperate with Ero1 to form disulfides ^23–27^. Incubation of isolated Pdi1 with Ero1 (for purity of Ero1, see **Extended Data Fig. 7a**) and RBun in the presence of molecular oxygen resulted in the formation of active RB, as shown by RNase activity with cCMP as the substrate (**Fig. 7a**). In contrast, Pdi1 in the Mnl1-Pdi1 complex was inactive, even when Ero1 was present in excess over Mnl1, at a molar ratio similar to the situation in *S. cerevisiae* cells ^52^ (**Fig. 7b**). Disulfide formation was observed with oxidized glutathione (GSSG) as the oxidant (**Fig. 7a**), but GSSG is not used *in vivo* and is in fact generated from reduced glutathione (GSH) entering the ER from the cytosol ^23,53^. The lack of Ero1-mediated activity is explained by Mnl1 blocking Ero1 binding to Pdi1; severe steric clashes were observed when our Mnl1-Pdi1 structure was compared with that of the Alphafold- predicted Ero1-Pdi1 complex (**Fig. 7c**). In addition, Mnl1 interacts with Pdi1 stronger than Ero1, as the Mnl1-Pdi1 complex is stable in gel filtration, whereas the Ero1- Pdi1 complex is not ^54^. Mnl1 binding would also interfere with face-to-face dimerization of Pdi1, which is required for the oxidative function of the human homolog ^55^.

**Figure 7.**
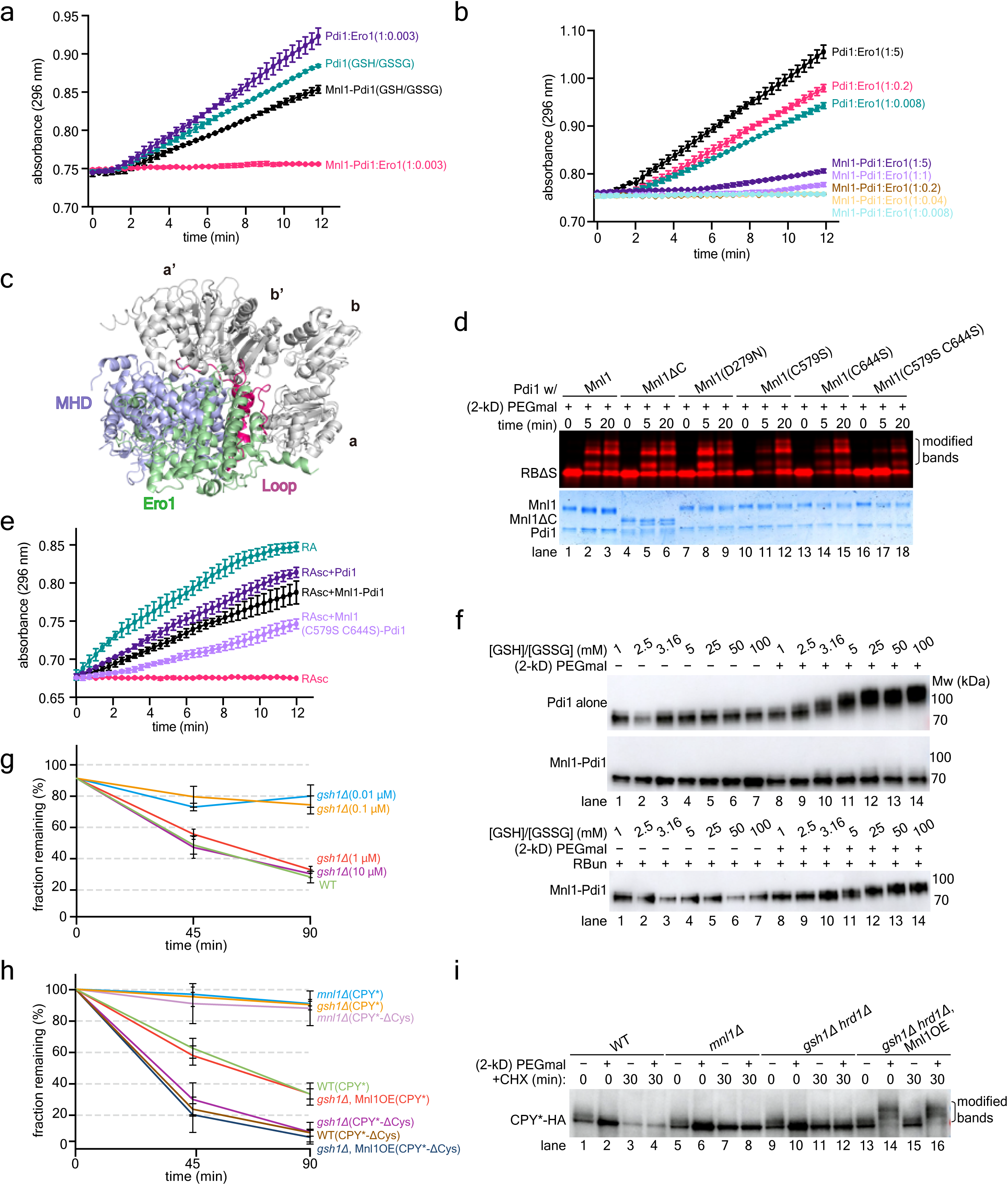
Mnl1-Pdi1 complex functions as a disulfide reductase in ERAD. **(a)** Unfolded RNase B (RBun) was incubated with isolated Pdi1 or Mnl1-Pdi1 complex (both 1.2 μM) and purified Ero1 (4 nM) in the absence of a redox buffer, so that molecular oxygen served as the oxidant. The renaturation of RNase B was followed by measuring the cleavage of cCMP over time. In parallel, reactions were performed in the absence of Ero1 with a redox buffer containing GSSG as the oxidant. Each point on the curves shows the means and standard deviation of three experiments. **(b)** As in (a), but with different concentrations of Ero1 (ranging from 9.6 nM to 6 μM). **(c)** Overlay of the structures of the Mnl1-Pdi complex and of the Alphafold-predicted Ero1-Pdi1 complex, based on the Trx domains of Pdi1. The model for Ero1 lacked the signal sequence and trans-membrane segment, and the CTD of Mnl1 was deleted for clarity. Note the extensive clashes. **(d)** RNase B lacking the S-peptide (RBΔS) was fluorescently labeled and incubated with wild-type or mutant Mnl1-Pdi1 complex in the presence of a redox buffer containing GSH. The reduction of disulfides was monitored by modification of the free cysteines with 2-kDa PEGmal. The samples were analyzed by SDS-PAGE, followed by fluorescence scanning and Coomassie blue staining. **(e)** The renaturation of scrambled RNase A (RAsc) was tested in a redox buffer containing GSH with isolated Pdi1, the Mnl1-Pdi1 complex, or a complex in which both Mnl1 loop cysteines were mutated. Controls were performed with only folded RA or RAsc. RNase activity was measured by following the cleavage of cCMP over time. Each point on the curves shows the means and standard deviation of three experiments. **(f)** Isolated Pdi1 or Mnl1-Pdi1 complex was incubated with 0.1 mM GSSG and increasing concentrations of GSH (0.1 to 10 mM). In the bottom panel, RBun was added to the Mnl1-Pdi1 complex. RBun was pretreated with iodoacetamide and the modification reagent removed by a desalting column. All samples were treated with TCA, the proteins dissolved in SDS, incubated with 2-kDa PEGmal, and analyzed by SDS-PAGE and immunoblotting for Pdi1. **(g)** ERAD of CPY*-HA was determined by cycloheximide (CHX) chase experiments in wild-type (WT) cells or cells lacking Gsh1 (*gsh1*Δ). The cells were grown in the presence of different concentrations of GSH (in brackets) before addition of CHX. The samples were analyzed by SDS-PAGE and immunoblotting for HA. The intensities of the CPY*-HA bands were quantified. Shown are the fractions of CPY*-HA remaining at different time points (means and standard deviation of three experiments). **(h)** As in (g), but following ERAD of CPY*-HA and a mutant lacking all cysteines (CPY*ΔCys) in the indicated cells grown in a low concentration of GSH (0.1 μM). Where indicated, Mnl1 was overexpressed from the GAL1 promoter (Mnl1OE). **(i)** As in (h), but only with CPY*-HA, and the samples were incubated with 2-kDa PEGmal to modify free thiols.

Pdi1 in the Mnl1-Pdi1 complex can still function as a disulfide reductase. When the Mnl1- Pdi1 complex was incubated with fluorescently labeled RBΔS in the presence of GSH, disulfide bonds were converted into free thiols, as shown by their modification with 2-kDa PEG-maleimide (PEGmal) (**Fig. 7d**, lanes 1-3). Abolishing the mannosidase activity of Mnl1 (lanes 7-9), or mutating one of the two cysteines in the Mnl1 loop had no effect (lanes 10-12 and 13-15), but mutating both cysteines moderately decreased the efficiency of disulfide reduction (lanes 16-18). The CTD was not required for disulfide reduction (lanes 4-6), even though this domain strongly promoted binding of RBΔS to the Mnl1-Pdi1 complex (**Fig. 4c**). As expected, the appearance of thiol-modified protein was dependent on the presence of Mnl1-Pdi1, GSH, and PEGmal, and did not occur with native RB (**Extended Data Fig. 7b**). Ero1 had no effect, even when added in excess (**Extended Data Fig. 7c**), suggesting that the Mnl1-Pdi1 complex can function as a reductase under the oxidizing conditions prevailing in the ER. Similar results were obtained with insulin as the substrate for disulfide bond reduction (**Extended Data Fig. 7d,e**). The efficiency of reduction was about equally efficient with Mnl1-Pdi1 and free Pdi1 (**Extended Data Fig. 7e**, lanes 10-12 versus 16-18).

The isomerase activity of Pdi1, a reaction that requires the transient reduction of disulfide bonds, was also not affected by the association with Mnl1. RAsc containing scrambled disulfides did not show RNase activity, but the activity was restored after incubation with either free Pdi1 or Mnl1-Pdi1 complex in a redox buffer containing GSH (**Fig. 7e**). The double cysteine mutant showed a partial defect (**Fig. 7e**). Again, Ero1 did not affect the reaction (**Extended Data Fig. 7f**). Taken together, these results show that Pdi1 can perform disulfide bond reduction when in complex with Mnl1.

To test the effect of Mnl1 on the redox behavior of Pdi1, we incubated isolated Pdi1 or Mnl1- Pdi1 complex with redox buffers containing increasing concentrations of GSH. The samples were precipitated with trichloroacetic acid, dissolved in SDS, and treated with 2-kDa PEGmal to modify free cysteines (**Fig. 7f**). As expected ^56^, isolated Pdi1 showed a large size shift at sufficiently high concentrations of GSH, indicating that free cysteines had been generated that could be modified by PEGmal. In contrast, no major size shift was observed with the Mnl1-Pdi1 complex (**Fig. 7f**), suggesting that Pdi1 remains in its oxidized state. However, after addition of RBun, in which all cysteines had been blocked by modification with iodoacetamide, the reduction of Pdi1 by GSH was partially restored (**Fig. 7f**). These results lead to a model in which substrate binding allows GSH to reduce Pdi1, which could then transfer the electrons to the substrate; after substrate dissociation, Pdi1 would revert back to the more stable, oxidized state in the Mnl1-Pdi1 complex.

### The Mnl1-Pdi1 complex reduces disulfide bonds of an ERAD substrate *in vivo*

Finally, we investigated whether the Mnl1-Pdi1 complex reduces disulfide bonds of ERAD substrates in *S. cerevisiae* cells. To this end, we followed the degradation of CPY*-HA, a protein with multiple disulfide bonds. Because GSH is the main disulfide reductant in the ER lumen, we employed cells lacking the glutathione-synthesizing enzyme Gsh1 ^23,53^. In this strain, the folding of wild-type CPY is normal ^23^. However, little degradation of CPY*-HA was observed in *gsh1*Δ cells when low concentrations of GSH were added (**Fig. 7g** and **Supplementary Fig. 2a**). The inhibition of ERAD was as strong as in the absence of Mnl1 (**Fig. 7h** and **Supplementary Fig. 2b**). At higher GSH concentrations, ERAD was restored (**Fig. 7g**), indicating that the reduction of disulfides is required for the degradation of CPY*-HA. Importantly, ERAD was also restored when the concentration of the Mnl1-Pdi1 complex was increased by overexpressing Mnl1 from a GAL1 promoter (**Fig. 7h**). Thus, the Mnl1-Pdi1 complex is required for the reduction of the disulfides of CPY*-HA. Consistent with this conclusion, a CPY* mutant lacking all cysteines (CPY*ΔCys) was degraded even faster than CPY* in wild-type cells (**Fig. 7h**), as this substrate does not need disulfide reduction. More importantly, the degradation of CPY*ΔCys was still dependent on Mnl1, as the mannosidase activity is required, but the degradation was no longer affected by the absence of GSH (**Fig. 7h**). Our results are consistent with data in the literature showing that Mnl1 function is sensitive to the cysteines in CPY* ^17^. Surprisingly, Mnl1 deletion has little effect in an *alg3*Δ mutant, in which a Man5 glycan species with a terminal α1,6 mannose-residue is transferred directly to CPY* ^8,57^, perhaps because CPY* does not form disulfide bonds in this strain.

To directly monitor the presence of free thiols in CPY*-HA, we used modification with 2- kDa PEGmal (**Fig. 7i**). In wild-type cells, CPY*-HA was barely modified, indicating that the cysteines were mostly engaged in disulfide bonds, both before addition of cycloheximide and after a 30 min chase (lanes 1-4). As expected, much of the protein was degraded during the 30 min chase (lanes 3 and 4 versus 1 and 2). To prevent degradation, we analyzed CPY*-HA in cells lacking ERAD components. In the absence of Mnl1 or of Gsh1 and Hrd1, the cysteines of CPY*- HA were again largely in the oxidized state before and after the chase (lanes 5-12). However, when Mnl1 was overexpressed, CPY*-HA was modified by PEGmal (lanes 13-16), indicating that the cysteines were no longer disulfide bonded. Thus, the Mnl1-Pdi1 complex reduces the disulfides of CPY*-HA in preparation of its retro-translocation into the cytosol.

## Discussion

Here, we show that the Mnl1-Pdi1 complex initiates ERAD-L by performing two crucial reactions, trimming of the glycan and reduction of disulfide bonds. The mannosidase Mnl1 in the complex uses its CTD to act on globular, misfolded proteins, generating an exposed α1,6- mannose residue that serves as a signal for degradation (**Fig. 8**). In a subsequent reaction, the Pdi1 component of the complex utilizes GSH to reduce the disulfides of the de-mannosylated protein, generating an unfolded polypeptide that can be retro-translocated across the ER membrane.

**Figure 8.** Model for the initiation of ERAD. The N-glycan of glycoproteins is trimmed by glycosidases; the three glucose residues (brown triangles) and one mannose (green circles) are removed to generate a Man8 species. At the same time, disulfide bridges are formed (red lines). If the protein does not reach its native folded state, a mannose residue is removed by the Mnl1 component of the Mnl1-Pdi1 complex, generating an exposed α1,6-mannose signal (purple circle). In the next step, the Pdi1 component of the Mnl1-Pdi1 complex uses GSH to reduce the disulfides of the substrate. The resulting unfolded polypeptide binds to the Hrd1 complex through the α1,6-mannose residue and an adjacent unstructured region. The protein is subsequently retro-translocated into the cytosol for degradation by the proteasome.

After N-glycosylation, all glycoproteins undergo initial glycan processing steps in the ER lumen, i.e. the removal of the three glucose residues by glucosidases and a mannose residue by Mns1, to generate the Man8 species (**Fig. 1a**). At the same time, many proteins form disulfide bonds in reactions that are catalyzed by Pdi1 and Ero1 or by other redox enzymes in the ER lumen. As a result, a protein that cannot reach its native folded state, is generally not entirely unfolded and instead adopts a globular structure (**Fig. 8**). Such misfolded, globular proteins are the preferred substrates for the mannosidase Mnl1, which generates the Man7 glycan with an exposed α1,6-mannose residue and irreversibly commits misfolded glycoproteins to ERAD. Our results show that Mnl1 ignores completely unfolded proteins that have not yet undergone folding attempts and need to be spared from degradation. However, if the products of the mannosidase reaction contain disulfide bonds, they cannot directly be retro-translocated across the ER membrane; the disulfides need to be reduced by Mnl1-associated Pdi1 and GSH to generate unfolded polypeptides with free cysteines. The unfolded polypeptides can then bind to the Hrd1 complex, employing both the exposed α1,6-mannose residue and an adjacent unstructured polypeptide segment, which bind to the Yos9 and Hrd3 components of the Hrd1 complex, respectively, thus initiating retro-translocation into the cytosol. The cascade of two quality control steps, one mediated by Mnl1 and the other by the Hrd1 complex, ensures that only terminally misfolded proteins are degraded, whereas folding intermediates are ignored.

Our results show that each component of the Mnl1-Pdi1 complex modifies the behavior of the other. Pdi1 changes the substrate specificity of Mnl1’s CTD, so that it switches from binding completely unfolded polypeptides to binding misfolded, globular proteins. Pdi1 also keeps Mnl1 soluble in the ER. *Vice versa*, Mnl1 blocks the interaction of Pdi1 with Ero1, so that Pdi1 cannot act as an oxidase. Instead, Mnl1 causes Pdi1 to function as the elusive disulfide reductase in ERAD. Unfolded polypeptide segments bind preferentially to Pdi1, which outcompetes the CTD. The binding of an unfolded polypeptide chain causes Pdi1 to detach at least partially from Mnl1. A dissociated redox-active Trx domain could then utilize GSH to reduce disulfides in substrates. After substrate release, Pdi1 would return to the more stable oxidized state that is enforced by its interaction with Mnl1.

Isolated Pdi1 can act as a net reductase *in vitro* (e.g. **Extended Fig. 7e**) and functions as a disulfide isomerase *in vivo*. In intact cells, only a small fraction of free Pdi1 might be in the reduced state required for disulfide reduction, whereas the majority of Pdi1 would be in the oxidized state required for disulfide bond formation. If free Pdi1 can act as a reductase *in vivo*, it is probably less efficient than Pdi1 in complex with Mnl1. Mnl1 could enhance disulfide reduction by its observed effect on the redox behavior of Pdi1 (**Fig. 7f**), or the flexible CTD could present globular misfolded proteins to the neighboring Pdi1 molecule for disulfide reduction *in vivo*. Such a hand-off might occur even if a substrate molecule transiently dissociates from the CTD after the mannosidase reaction.

The function of the Mnl1-Pdi1 complex is likely conserved in all eukaryotes. The Mnl1 homolog in *S. pombe* contains two domains following the MHD (**Extended Data Fig. 7g**), with the intermediate domain structurally similar to the CTD of *S. cerevisiae* Mnl1 (not shown). EDEM3 is probably the mammalian homolog of Mnl1, but its two C-terminal domains are unrelated to the CTD of *S. cerevisiae* Mnl1 (**Extended Data Fig. 7g**). The first of these domains has a predicted hydrophobic groove that might bind substrate. Mammals have two other EDEM homologs (EDEM1 and EDEM2), which contain essentially only a mannosidase domain (**Extended Data Fig. 7g**); EDEM2 may be a functional homolog of yeast Mns1 ^58,59^. All three EDEM proteins have been reported to associate with PDI-like enzymes ^33,59,60^, but it is unclear whether they form stable, stoichiometric complexes with their redox partners and whether these partners serve as disulfide reductases in ERAD, similar to the EDEM1 partner ERdj5 ^33,34^.

Other ER proteins that stably interact with PDI or its homolog ERp57 (prolyl 4-hydroxylase, MTP, and tapasin) also interact with unfolded (poly)peptides (collagen, apolipoprotein, antigenic peptides). In the case of prolyl 4-hydroxylase, hydroxylation of prolines is followed by the assemply of pro-collagen into a triple helix, which starts with the formation of disulfides in a C-terminal domain and determine which collagen chains associate with one another (a “cysteine code”) ^61^. The cysteines need to be in a reduced state before assembly of the triple helix, raising the intriguing possibility that PDI in the complex might function as a reductase, as in the Mnl1 complex.

## Acknowledgements

We thank Kaiku Uegaki for helpful discussions, Michael Skowyra for help with the bead binding assay, Haipeng Guan for help with cryo-EM data analysis, and Kaiku Uegaki, Rudolf Pisa, Vlad Denic, and Pedro Carvalho for critical reading of the manuscript. We thank the Core for Imaging Technology & Education (CITE) and the Center for Macromolecular Interactions at Harvard Medical School for help with experiments. We thank S. Sterling, R. Walsh, and M. Mayer at the Harvard Cryo-EM Center for Structural Biology for help in microscope operation and data collection. This work was supported by a NIGMS grant (R01 GM052586) to T.A.R. T.A.R. is a Howard Hughes Medical Institute Investigator.

## Author contributions

D.Z. performed all biochemical and cellular experiments, and performed cryo-EM data analysis on one particle class. X.W. designed the purification scheme for the Mnl1-Pdi1, determined the cryo-EM structure from one particle class, and designed the mannosidase assay. T.A.R. supervised the project. T.A.R. and D.Z. wrote the manuscript with input from X.W.

## Ethics declarations

The authors declare no competing interests.

## Methods

### Yeast strains and cultures

*S. cerevisiae* strain INVSc1 was obtained from Thermo Fisher Scientific. BY4741, BY4743, *mnl1*Δ*, gsh1*Δ *, ubc7*Δ and *pep4*Δ were obtained from Horizon Discovery. Strains with multiple gene deletions were constructed by PCR-based homologous recombination. Plasmids encoding *S. cerevisiae* proteins were transformed into wild-type *S. cerevisiae* cells or strains lacking the indicated genes. Transformed yeast cells were grown for 3 days on synthetic amino acid dropout (SD) plates. For protein purifications, yeast colonies were picked and cultured for 24 h in minimum medium at 30°C. The starter culture was diluted 1:50 and grown at 30°C for 24 h. Protein expression was induced with YPG (1% yeast extract, 2% bacto-peptone and 2% galactose). The cells were harvested after incubation at 27°C for 18 h.

### Mammalian cell cultures

FreeStyle^TM^ 293-F (Thermo Fisher) cells were cultured in FreeStyle^TM^ 293 expression medium supplemented with Fetal Bovine Serum (Thermo Fisher) at 37 °C for 2 to 3 days and diluted twice before transfection. Plasmids coding for MRH-His6-IgM (FC region), MBP-Mnl1-CTD-His6, MBP- Mnl1-mCTD-His6, or MBP-His6 were transfected into FreeStyle^TM^ 293-F cells. For 1 L HEK293 culture, 1 mg of plasmid was incubated with 3 mg of Linear PEI 25K (Polysciences) in 100 mL of Opti-MEM (Thermo Fisher) medium at room temperature for 25 min. The mixture was added dropwise into the medium containing HEK293 cells at a density of 2-2.5 million/mL. The cells were cultured at 37 °C for 16 h before addition of 10 mM sodium butyrate to boost expression. The medium containing secreted protein was harvested 48 h post transfection.

### Plasmids

The Mnl1-Pdi1 complex was expressed from a modified version of the pRS42X vector (pRS42X- LNK) ^62^. This plasmid allows the insertion of multiple expression cassettes into the same vector. Both Mnl1 and Pdi1 were expressed under the GAL1 promoter. Pdi1 was untagged while Mnl1 had a FLAG tag at its C-terminus. Pdi1 with a C-terminal SBP tag, Mns1 (amino acids 26-549) with a N-terminal His14 tag and a C-terminal FLAG tag, and Ero1 (amino acids 1-424) with a C-terminal HA tag replacing its trans-membrane segment were also expressed under the GAL1 promoter from the pRS42X vector. CPY* was expressed under the GAL1 promoter from the pRS42X vector. The protein contained an N-terminal His14 tag and a C-terminal SBP tag, followed by the ER retention signal HDEL. CPY* with a C-terminal HA tag was cloned into the pRS31X vector and expressed under its native promoter. Plasmids expressing Mnl1 or Mnl1 mutants were cloned into the pRS41X vector and expressed under the native Mnl1 promoter.

The MRH domain of mammalian OS9 was fused to the constant region (FC) of IgM and cloned into the pCAGEN vector. The fusion protein contained the signal sequence of human IgG κ-light chain at the N-terminus. MBP alone, MBP fused to the CTD, and MBP fused to the mCTD were also cloned into the pCAGEN vector.

### Immunoblotting and antibodies

Antibodies used in this study were: anti-FLAG antibody from rabbits (Millipore, 1:3000), anti-HA antibody from rats (Millipore, 1:2000), anti-PGK1 antibody from mice (Abcam, 1:3000), anti-PDI antibody from mice (Thermo Fisher, 1: 2000), anti-MBP antibody from mice (New England Biolabs, 1:3000), Goat anti-mouse IgG HRP conjugated (Thermo Fisher, 1: 3000), Goat anti-rabbit IgG HRP conjugated (Thermo Fisher, 1: 3000), Goat anti-rat IgG HRP conjugated (Thermo Fisher, 1: 3000).

### Purification of proteins expressed in *S. cerevisiae*

For purification of the Mnl1-Pdi1 complex, 100 g of cell pellet were resuspended in buffer A (25 mM HEPES pH 7.4, 150 mM NaCl) supplemented with 2 mM phenylmethanesulfonyl fluoride (PMSF) and 2 mM pepstatin A. The cells were lysed in a BioSpec BeadBeater for 45 min with 20 s/60 s on/off cycles in a water-ice bath. Cell debris were pelleted by centrifugation at 5,000 g for 20 min. The supernatant was subjected to centrifugation in a Beckman Ti45 rotor at 43,000 rpm for 1.5 h at 4°C. The pelleted membranes were resuspended with a Dounce homogenizer in buffer A. The membranes were solubilized in 200 mL buffer A containing 1.5% Triton-X100 and a protease inhibitor cocktail for 1 h at 4°C. Insoluble material was removed by centrifugation in a Beckman Ti45 rotor at 43,000 rpm for 40 min. The supernatant was incubated with 2 mL anti- FLAG M2 resin for 3 h. The beads were washed with 20 mL of buffer A containing 1% Triton-X and then with 30 mL of buffer A lacking detergent. The proteins were eluted with buffer B (25mM HEPES pH7.4, 300 mM NaCl, 5% glycerol) supplemented with 3x FLAG peptide (Sigma). The eluted material was applied to a Superdex 200 Increase 10/300GL Increase column, equilibrated with buffer A. Peak fractions were pooled and concentrated to 3-4 mg/mL for cryo- EM analysis. All Mnl1 mutants and Mns1 were purified similarly.

Pdi1-SBP was purified from a detergent-solubilized membrane extract by incubating with streptavidin agarose resin for 2 h. The beads were washed with 10 column volumes of buffer A containing 1% Triton-X and 15 column volumes of buffer A. The protein was eluted with buffer A supplemented with 2 mM biotin, and was applied to a Superdex 200 Increase 10/300GL Increase column equilibrated with buffer A. Ero1(1-424)-HA was purified similarly with anti-HA resin, and the protein was eluted with anti-HA peptide and was applied to a Superdex 200 Increase 10/300GL Increase column equilibrated with buffer A.

For purification of His14-CPY*-SBP-HDEL, 150 g of cell pellet were resuspended in buffer C (25 mM HEPES pH 7.4, 300 mM NaCl, 25 mM imidazole). The cells were lysed and the membranes were collected. The membranes were resuspended in buffer D (25 mM HEPES pH 7.4, 500 mM NaCl, 8 M urea, 25 mM imidazole) for 1 h to release luminal proteins. The membranes were pelleted and the supernatant incubated with 5 mL Ni-NTA resin for 2 h at 4°C. The beads were sequentially washed with buffer E (25 mM HEPES pH 7.4, 500 mM NaCl, 6 M urea, 25mM imidazole), buffer F (25 mM HEPES pH 7.4, 500 mM NaCl, 2 M urea, 25 mM imidazole), and buffer G (25 mM HEPES pH 7.4, 500 mM NaCl, 80 mM imidazole). The protein was eluted with buffer H (25 mM HEPES pH 7.4, 500 mM NaCl, 500 mM imidazole, 1 mM DTT). The eluted material was incubated with GST-3C protease overnight to cleave off the His tag, and further incubated with 1 mL streptavidin agarose resin for 1 h. The beads were washed with buffer I (25 mM HEPES pH 7.4, 500 mM NaCl, 1 mM DTT) and protein eluted with buffer J (25 mM HEPES pH7.4, 500 mM NaCl, 10% glycerol, 2 mM biotin, 1 mM DTT). The eluted material was applied to a desalting column (Thermo Scientific) equilibrated with buffer K (25 mM HEPES pH 7.4, 500 mM NaCl, 10% glycerol, 1 mM DTT).

### Purification of proteins expressed in FreeStyle^TM^ 293-F cells

The purification of the His-tagged proteins was carried out as described ^63^, with some modifications. The collected medium was supplemented with 50 mM Tris pH 8.0, 200 mM NaCl, 20 mM imidazole, 1 μM NiSO_4_ and incubated with Ni-NTA beads. The beads were washed extensively with 25 mM HEPES pH 7.4, 200 mM NaCl, 20 mM imidazole. Protein was eluted with 25 mM HEPES pH 7.4,200 mM NaCl, 300 mM imidazole. Eluted MRH-His6-IgM was applied to a Superdex 200 Increase 10/300GL Increase column equilibrated with 25 mM HEPES pH 7.4, 150 mM NaCl, 5% glycerol. For MBP fusions, the eluted protein was concentrated and buffer exchanged into 25 mM HEPES pH 7.4, 150 mM NaCl, 5% glycerol.

### Cryo-EM sample preparation and data acquisition

3 μL of the Mnl1-Pdi1 complex at 1mg/mL was applied to a glow-discharged Quantifoil grid (1.2/1.3, 400 mesh). The grids were then blotted for 7.5 s at ∼90 % humidity and plunge-frozen in liquid ethane using a Vitrobot Mark IV (Thermo Fisher Scientific).

Cryo-EM data were collected on a Titan Krios electron microscope (Thermo Fisher Scientific) operated at 300 kV and equipped with a K3 Summit direct electron detector (Gatan) at Harvard Cryo-EM Center for Structural Biology. A Gatan Imaging filter with a slit width of 20 eV was used. All cryo-EM movies were recorded in counting mode using SerialEM. The nominal magnification of 105,000x corresponds to a calibrated pixel size of 0.83 Å on the specimen. The dose rate was 20 electrons/Å^2^/s. The total exposure time was 3.5 s, resulting in a total dose of 70.3 electrons/Å^2^, fractionated into 60 frames (59 ms per frame). The defocus range for was between 0.8 and 2.2 μm. All parameters of EM data collection are listed in Table S1.

### Image processing

Dose-fractionated super-resolution movies were subjected to motion correction using the program MotionCor2 ^64^, with dose-weighting. The program CTFFIND4 (ref. ^65^) was used to estimate defocus values of the summed images from all movie frames. Particles were autopicked by crYOLO ^66^. After manual inspection to discard poor images, 2D and 3D classifications were done in Relion 3.0 (ref. ^67^). 2,413,957 picked particle images were extracted and subjected to two rounds of 2D classifications to remove junk particles, which resulted in 1,971,531 particles. After one round of global 3D classification using an initial model generated by Relion 3.0, 1,086,314 particles from one class with good protein features were selected for 3D refinement. After a second round of 3D classification, 313,324 particles from one class with more complete structural features (complete) were selected and then subjected to 3D refinement using a mask surrounding the protein, followed by particle polishing and CTF refinement. Polished particles were subjected to another round of 3D refinement. 443,216 particles from another class with the CTD of Mnl1 missing (incomplete) were processed in a similar way, following particle polishing, CTF refinement and 3D refinement.

Local resolutions were calculated, and map sharpening was performed in Relion 3.0. All reported resolutions are based on gold-standard refinement procedures and the FSC=0.143 criterion.

### Model building

The model for Mnl1-Pdi1 complex was built using an Alphafold model of Mnl1 and the crystal structure of Pdi1 (PDB code: 2B5E) as initial models. For Pdi1, the four Trx-like domains were in a slightly different arrangement than the crystal structure. Each Trx-like domain was first fitted into its corresponding cryo-EM densities as a rigid body and then manually modified and connected according to the density. Amino acids 24-500 of Pdi1 could be modeled. For Mnl1, the Alphafold models of the MHD and CTD were first placed into the cryo-EM densities as rigid bodies and then modified manually according to the density. Then, the long loop between MHD and CTD domain was built *de novo* according to the density map. The model of the Mnl1-Pdi1 complex was then refined in Phenix ^68^.

### RNase preparation

RBΔS was prepared as described ^18^, with some modifications. Approximately 0.5 mg RNase B (RB) was incubated with 10 μg subtilisin (Sigma) in 25 mM HEPES pH 7.4, 150 mM NaCl. After incubation at 4°C for 16 h, 10 μg subtilisin was added for another 2 h. The pH was adjusted to 2.0 with hydrochloric acid and the mixture was kept for 1 h on ice to inactivate subtilisin. 10% trichloroacetic acid (TCA) was added for 12 h to precipitate RBΔS, and the mixture was centrifuged at 12,000 g for 10 min. The supernatant was removed and the pellet was dissolved in 8 M urea. The TCA precipitation was repeated to remove residual S-peptide. RBΔS was then dissolved in urea, and buffer exchanged into 25 mM HEPES pH 7.4, 150 mM NaCl, 5% glycerol.

Reduced and denatured RBun were prepared by incubating 5 mg RB with 6 M guanidine hydrochloride and 100 mM DTT in Tris-acetate pH8.0 at 25 °C for 18 h. Immediately before use, RBun was dialyzed overnight in a 50 mL-Falcon tube with nitrogen gas added to avoid oxidation.The solution was then buffer exchanged on a desalting column (Thermo Scientific) to further remove guanidine hydrochloride and DTT. ^35^S-methionine was used to monitor the efficiency of the removal of small molecules (less than 0.01 % left). To prepare scrambled RNase A (RAsc), RNase A was first reduced, and then treated with 25 mM N,N, Nʹ,Nʹ-tetramethylazodicarboxamide (diamide) at 25 °C for 1 h. The solution was then buffer exchanged.

### Protein Labeling

MRH-IgM was incubated with a 2:1 molar excess of DyLight 680 or DyLight 800 NHS ester for 1 h on ice. CPY* was incubated with a 2:1 molar excess of DyLight 800 NHS ester. RNase B (RB), RBΔS, or RBun was incubated with a 2:1 molar excess of DyLight 680 NHS ester. Excess dye was removed by gel filtration or dye-removal columns (Thermo Scientific).

Labeling with biotin was performed with RB, RBΔS or RBun by incubating the proteins with a 20:1 molar excess of NHS-PEG_4_-Biotin for 2 h on ice. Excess of NHS-PEG_4_-Biotin was removed with a desalting column (Thermo Scientific).

### Mannosidase assays

Substrate was incubated with Mnl1-Pdi1 complex and Mns1 at a ratio of 2 μM: 0.2 μM: 0.15 μM in 25 mM HEPES pH 7.4, 150 mM NaCl, 0.1% Nonidet P-40, 3 mM GSH, 0.3 mM GSSG, 2 mM CaCl_2_ at 30°C for different time periods (0 min, 10 min, 30 min, 60 min, 120 min). 20 mM EDTA was added to inhibit the reaction. Samples were then applied to 5 μL streptavidin resin equilibrated in IP buffer (25 mM HEPES pH 7.4, 150 mM NaCl, 0.1% Nonidet P-40) at 4°C for 30 min. Beads were washed with 200 μL IP buffer three times. 2 μM MRH-IgM was added and samples were incubated at 4°C for 30 min. Beads were then washed with 200 μL IP buffer three times and proteins were eluted in 50 μL (25 mM HEPES pH 7.4, 150 mM NaCl, 1% SDS, 2 mM biotin). Samples were diluted with loading buffer and subjected to SDS-PAGE. Fluorescently labeled CPY* and MRH-IgM were detected by fluorescence scanning on an Odyssey imager (LI-COR). RB, RBΔS, and RBun were detected using Coomassie blue staining.

The fluorescence in bands was quantitated using the ImageStudio software (LI-COR). For each lane, a rectangular box was selected to determine the total intensity of a band. The box size was kept constant for all bands on the same gel. An additional box of the same size was drawn over an empty region to determine background intensity. Signal intensity of each band was calculated as (total intensity –background intensity). The numbers for MRH-IgM were divided by those for CPY*. The resulting ratios were normalized to that at time-point zero.

### Cycloheximide-chase degradation assays

Cycloheximide-chase experiments were performed as described ^15^, with some modifications. Mid-log phase cells (0.4 to 0.6 OD_600_/mL) cultured in 50 mL liquid media at 30°C were used. Cells were mixed with fresh medium supplemented with 100 mg/mL cycloheximide to generate a final density of 2 OD_600_/mL. 4 OD_600_ units of cells were harvested at the indicated time points. Cells were lysed by vortexing for 2 min with 250 μL of acid-washed glass beads (0.1mm, Bio-Spec) and 200 μL of lysis buffer (10 mM MOPS, pH 6.8, 1% SDS, 8 M urea, 10 mM EDTA, 1x protease inhibitors cocktail). 200 μL of urea-containing sample buffer (125 mM Tris pH 6.8, 4% SDS, 8 M urea, 100 mM DTT, 10% glycerol, bromophenol blue) was added. The samples were incubated at 65°C for 5 min, centrifuged at 12,000 rpm, and subjected to SDS-PAGE and immunoblotting. HA-tagged substrate was detected using anti-HA (Millipore). PGK was detected using anti-PGK antibodies (Abcam) and served as a loading control. For quantification, the immunoblots were scanned with an Image Quant 800 Western blot imaging system (Amersham) and the intensities of the CPY*-HA and PGK bands were determined with Fiji ImageJ. For each time point, the intensity of the CPY* band was divided by that of the PGK band. These numbers were converted into percentages, setting that at time point zero to 100%.

For experiments in which Mnl1 was expressed from the GAL1 promoter and glutathione was added, cells were grown for three doubling times in medium containing 2% raffinose and the indicated concentrations of glutathione. The cells were then incubated in medium containing 2% galactose and glutathione for 6 h before performing cycloheximide-chase experiments.

### Co-immunoprecipitation of Mnl1 and Pdi1

Approximately 50 OD_600_ units of cells were harvested and resuspended in IPB buffer (25 mM HEPES pH 7.4, 200 mM NaCl) supplemented with a protease inhibitor cocktail. Cells were lysed with glass beads and cell debris were removed by centrifugation at 6,000 g for 1 min. Membrane fractions were collected by centrifugation with a TLA55 rotor (Beckman) at 42,000 rpm for 20 min. Membranes were solubilized in IPB containing 1% Nonidet P-40 for 1 h. The supernatant was incubated with 7 μL of anti-FLAG M2 resin for 2 h. The beads were washed three times with IPB containing 0.1% Nonidet P-40, and proteins were eluted with this buffer supplemented with 3x FLAG peptide. Eluted proteins were subjected to SDS-PAGE and immunoblotting. FLAG-tagged Mnl1 and Pdi1 were detected using anti-FLAG (Millipore) and anti-Pdi1 (38H8, Thermo Fisher) antibodies, respectively.

To test for co-immunoprecipitation of Mnl1 and Pdi1 after reduction of disulfide bonds, cells were harvested and resuspended in IPB buffer supplemented with 10 mM DTT and a protease inhibitor cocktail. Cells were lysed and cell debris were removed by centrifugation. Membrane fractions were collected by centrifugation, washed to remove DTT, and solubilized in IPB containing 1% Nonidet P-40. The supernatant was incubated with anti-FLAG M2 resin, and the beads were collected and proteins were eluted.

### Pull-down experiments to detect substrate binding

DyLight 680 NHS ester-labeled RB, RBΔS or RBun was incubated with the Mnl1-Pdi1 complex at a ratio of 0.2 μM: 1 μM in a reaction buffer (25 mM HEPES pH 7.4, 150 mM NaCl, 0.1% Nonidet P-40, 3 mM GSH, 0.3 mM GSSG, 2 mM CaCl_2_) at 30°C for 30 min. In some experiments, S peptide or S peptide mutant (F8W H12A D14A) (synthesized by GenScript) was added. The mixture was incubated with 7 μL of anti-FLAG M2 resin for 1.5 h. The beads were washed three times with 25 mM HEPES pH 7.4, 150 mM NaCl, 0.1% Nonidet P-40, and proteins were eluted in this buffer supplemented with 3x FLAG peptide. Eluted proteins were subjected to SDS-PAGE. FLAG-tagged Mnl1 was detected using anti-FLAG antibody, and RB, RBΔS and RBun were detected by fluorescence scanning.

DyLight 680 NHS ester-labeled RB, RBΔS or RBun was also mixed with MBP-CTD at a ratio of 0.2 μM: 1 μM in a reaction buffer (25 mM HEPES pH 7.4, 150 mM NaCl, 0.1% Nonidet P-40). The mixture was incubated with anti-MBP magnetic beads (New England Biolabs) for 1 h. The beads were washed three times with the same buffer, and proteins were eluted in SDS buffer and subjected to SDS-PAGE.

### Testing protein aggregation by light scattering

300 nM luciferase or citrate synthase were incubated with purified MBP alone, MBP-CTD, or MBP-mCTD at different molar ratios (1:0.5, 1:1, 1:2, 1:3) in 50 mM Tris-HCl pH 7.5, 250 mM NaCl. Light scattering was measured with a DynaPro Plate Reader III (Wyatt Technologies) using discrete temperature increments (25°C-65°C). The hydrodynamic radius (R_h_) of particles and their relative intensity was measured. The relative intensity of particles (>200 nm) was quantitated.

### Substrate interference of disulfide crosslinking between Mnl1 and Pdi1

RBΔS, RBun, or RBsc were incubated with Mnl1-Pdi1 complex at the indicated molar ratios in a reaction buffer (25 mM HEPES pH 7.4, 150 mM NaCl, 0.1% Nonidet P-40, 2 mM CaCl_2_) at 30°C for 30 min. 2,2ʹ- dipyridyldisulfide (DPS) (Sigma) was added and the mixture was incubated at 30°C for 30 min. The reaction was terminated by addition of N-ethylmaleimide (NEM). The mixture was then subjected to SDS-PAGE and Coomassie blue staining.

### Substrate binding determined with bead-immobilized Mnl1-Pdi1 complex

Mnl1-Pdi1, Mnl1(C579S C644S)-Pdi1, or Mnl1ΔC-Pdi1 complex was incubated with DPS at 30°C for 2 h. Fluorescently labeled RB, RBun, or RBΔS was then added at a ratio of 4 μM: 2 μM. The mixture was incubated with 25 μL diluted (60x) anti-FLAG M2 resin in 25 mM HEPES pH 7.4, 150 mM NaCl in a 384-well glass-bottom plate (Cellvis) and kept at 25 °C for 2 h. Images were acquired with a spinning disk confocal microscope at Harvard Nikon Imaging Center. Fluorescence intensity was determined by measuring the intensity of circular regions (40 x 40 pixels) centered around the beads. The fluorescence of the surrounding was also determined, and six images were averaged. These numbers were used for flatfield correction to eliminate uneven illumination.

### RNase re-folding assays

The renaturation of RNase was followed by determining ribonuclease activity spectrophotometrically with cCMP as substrate. 4 mM cCMP was incubated with 1.2 μM of Mnl1-Pdi1 or the Mnl1(C579S C644S)-Pdi1 complex, or with Pdi1 alone in 100 mM Tris-acetate pH 8.0, 1 mM GSH, 0.2 mM GSSG. The assay was initiated by the addition of denatured or scrambled RNase (RBun or RAsc). The hydrolysis of cCMP was recorded continuously by following the absorbance at 296 nm. Where indicated, the GSH: GSSG buffer was replaced by Ero1.

### Determining the *in vivo* redox state of CPY*

Approximately 4 OD_600_ units of cells were harvested and suspended in 10% TCA. Cells were lysed with glass beads and collected by centrifugation. The pellets were washed three times with 100% cold acetone, and proteins solubilized and modified in 1% SDS, 100 mM sodium phosphate buffer pH 7.0, 150 mM NaCl, 4 M urea, 3 mM PEGmal (2-kDa) by incubation at 25°C for 2 h. The reaction was stopped by adding sample loading buffer containing 50 mM DTT, and the samples were subjected to SDS-PAGE and immunoblotting.

### Redox titrations with purified proteins

Pdi1 or Mnl1-Pdi1 complex (0.5 μM) was incubated in 100 mM sodium phosphate pH 7.0, 150 mM NaCl supplemented with 0.1 mM GSSG and various concentrations of GSH at 25°C for 1 h. The proteins were precipitated by incubation with 10% TCA on ice for 30 min, and the mixture was centrifuged at 15,000 g for 30 min. The pellet was washed twice with 100% cold acetone. The proteins were dissolved and modified in 1% SDS, 100 mM sodium phosphate pH 7.0, 150 mM NaCl, 3 mM PEGmal (2-kDa) by incubation at 25°C for 1 h. The reaction was stopped by adding sample loading buffer containing 50 mM DTT, and the samples were subjected to SDS-PAGE.

To test the effect of RBun in redox titration experiments with Mnl1-Pdi1, Rbun was treated with 40 mM iodoacetamide followed by removal of the reagent on a desalting column (Thermo Scientific). The Mnl1-Pdi1 complex was incubated with Rbun at a ratio of 0.2 μM: 4 μM in various GSSG: GSH buffers before sample processing as above.

### Test for *in vivo* disulfide bond formation between Mnl1 and Pdi1

Approximately 10 OD_600_ units of cells were harvested and resuspended in IPB buffer (25 mM HEPES pH 7.4, 200 mM NaCl) supplemented with a protease inhibitor cocktail and 100 mM iodoacetamide. Cells were lysed with glass beads and membrane fractions were collected. Membranes were solubilized in IPB containing 1% Nonidet P-40 and 100 mM iodoacetamide for 1 h, and the insoluble material was removed. The supernatant was incubated with 5 μL of anti- FLAG M2 resin for 2 h. The beads were washed three times with IPB containing 0.1% Nonidet P- 40, and proteins were eluted with this buffer supplemented with 3x FLAG peptide. Eluted proteins were subjected to SDS-PAGE and immunoblotting. FLAG-tagged Mnl1 and Pdi1 were detected using anti-FLAG (Millipore) and anti-Pdi1 (38H8, Thermo Fisher) antibodies, respectively.

## Data availability

The data supporting the findings of this study are available in the Electron Microscopy Bank and Protein Data Bank under accession codes EMD-60365 and PDB ID 8ZPW. The Alphafold model of Mnl1 and the crystal structure of Pdi1 (PDB 2B5E) were used for comparisons and as an initial model. Source data are provided with this paper.

## Notes

### Competing Interest Statement

The authors have declared no competing interest.

