## Supplementary figures and images for "Initiation of ERAD by the bifunctional complex of Mnl1 mannosidase and protein disulfide isomerase"

### ED figure 1

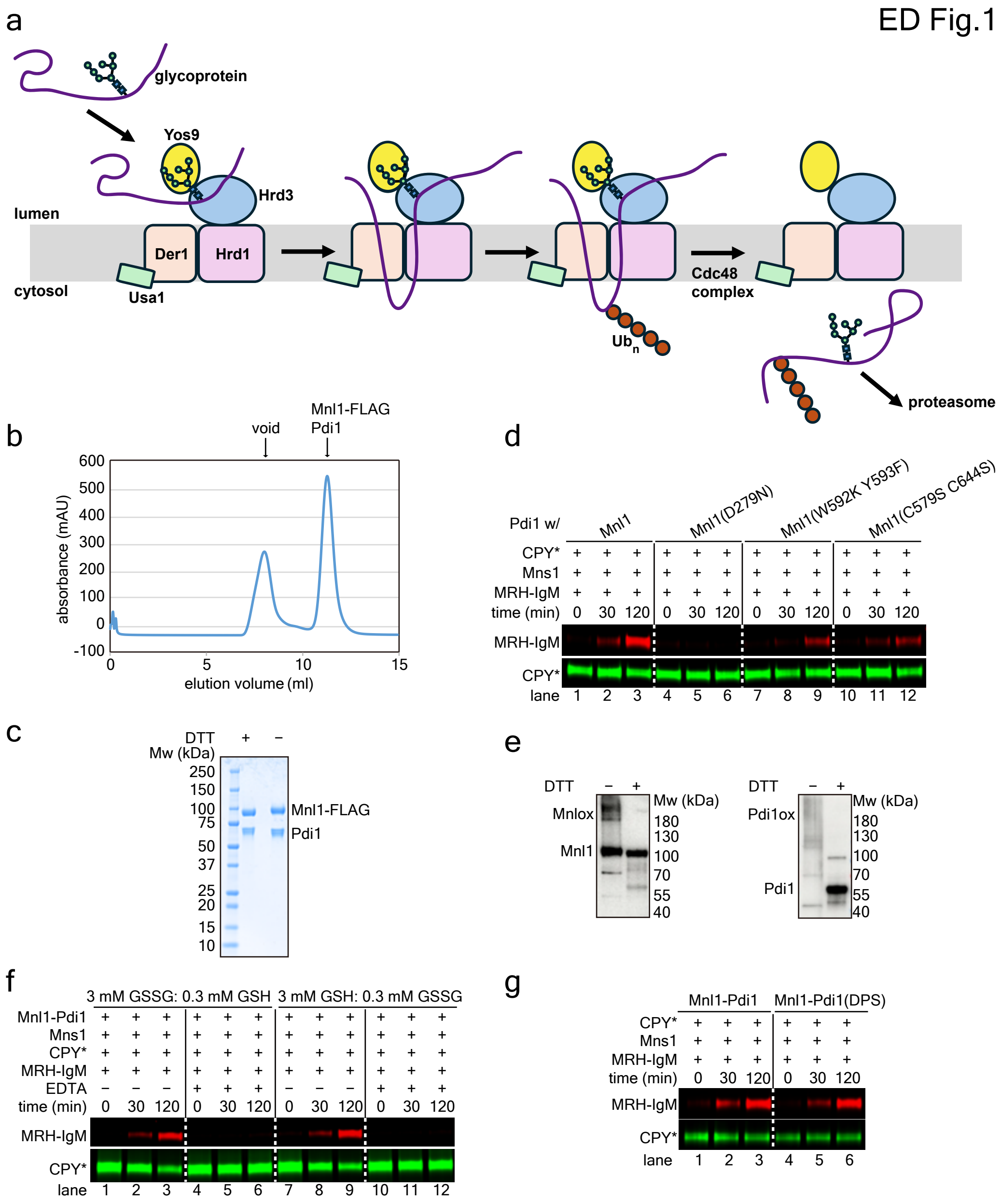

### ED figure 2

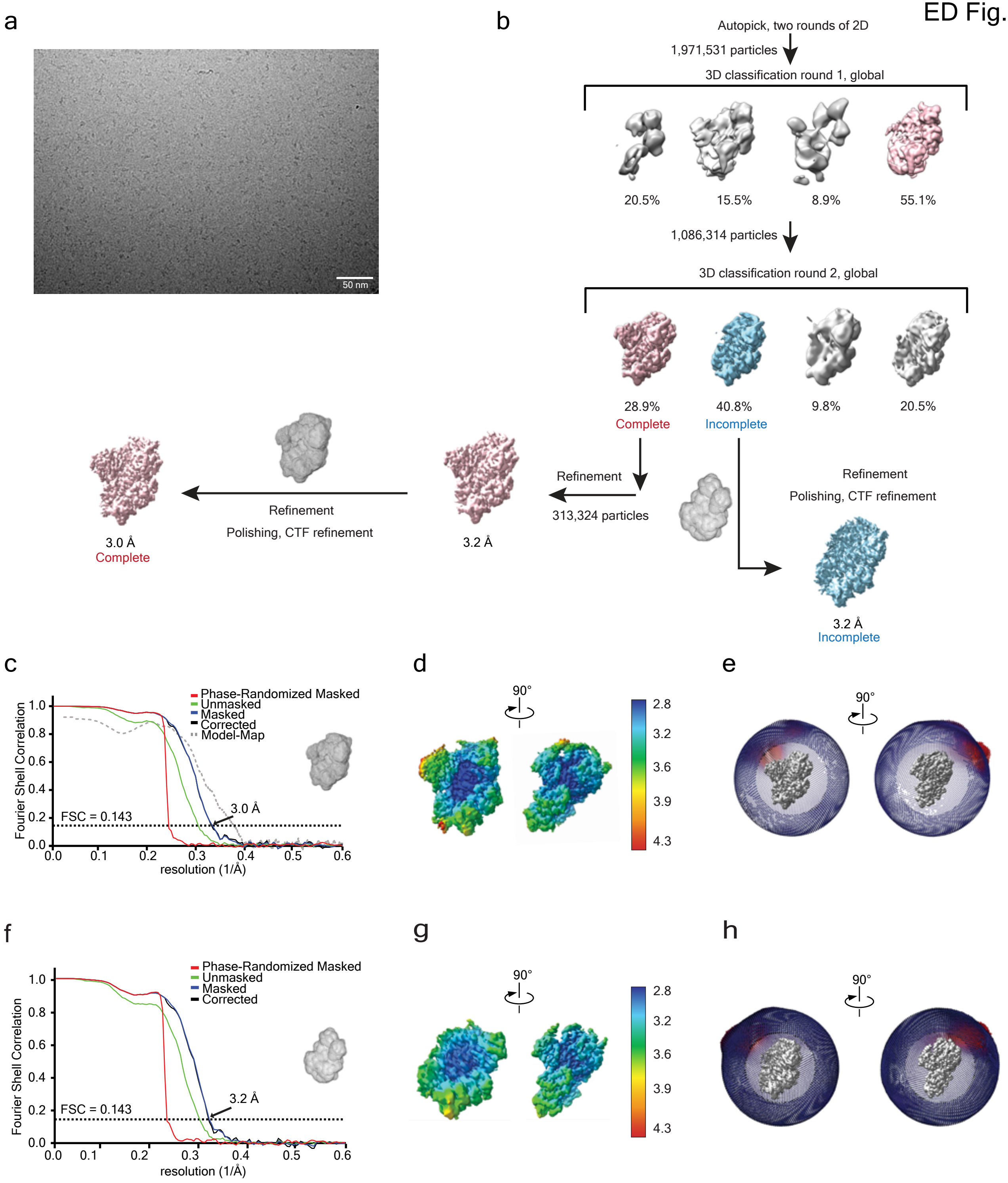

### ED figure 3

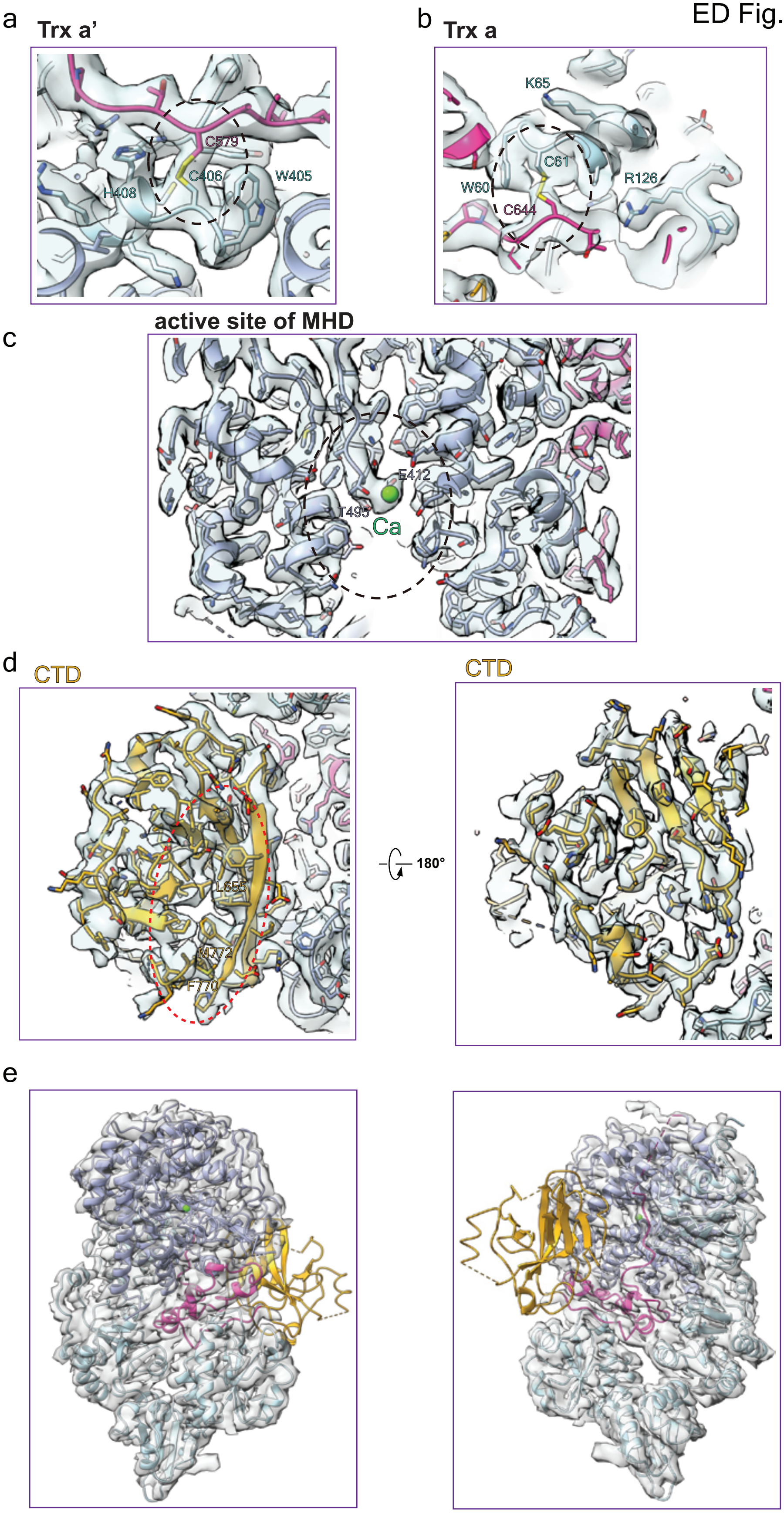

### ED figure 4

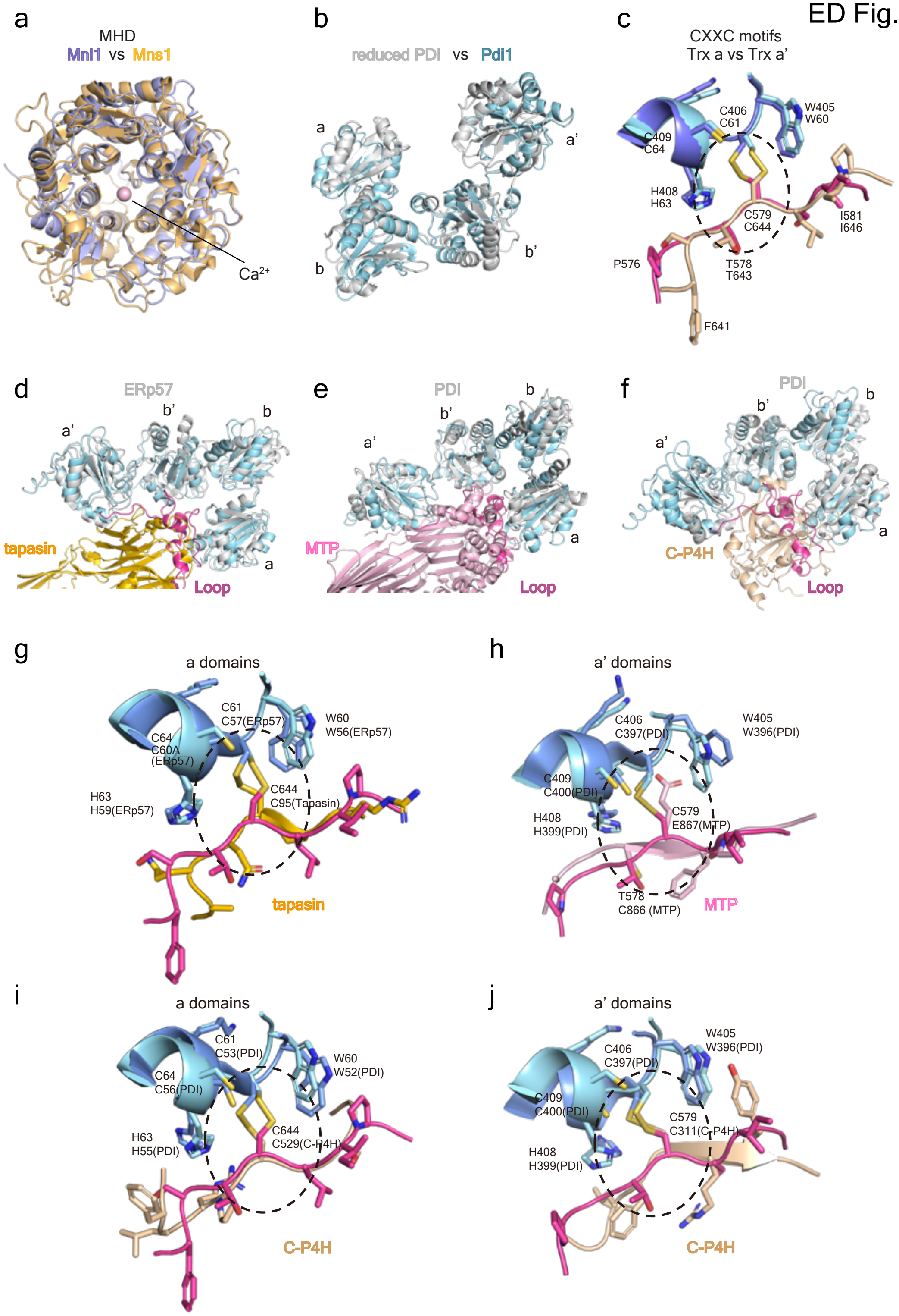

### ED figure 5

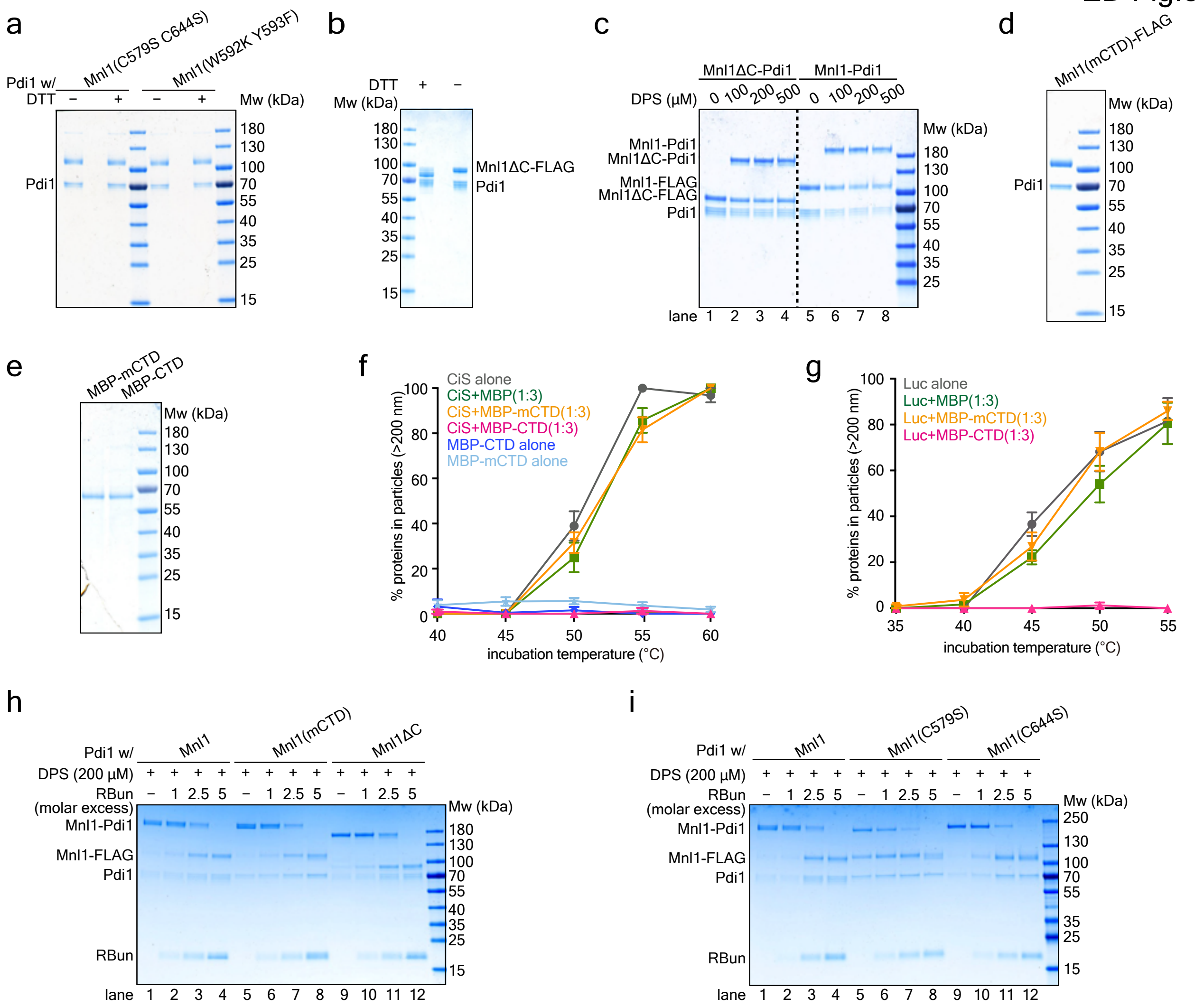

### ED figure 6

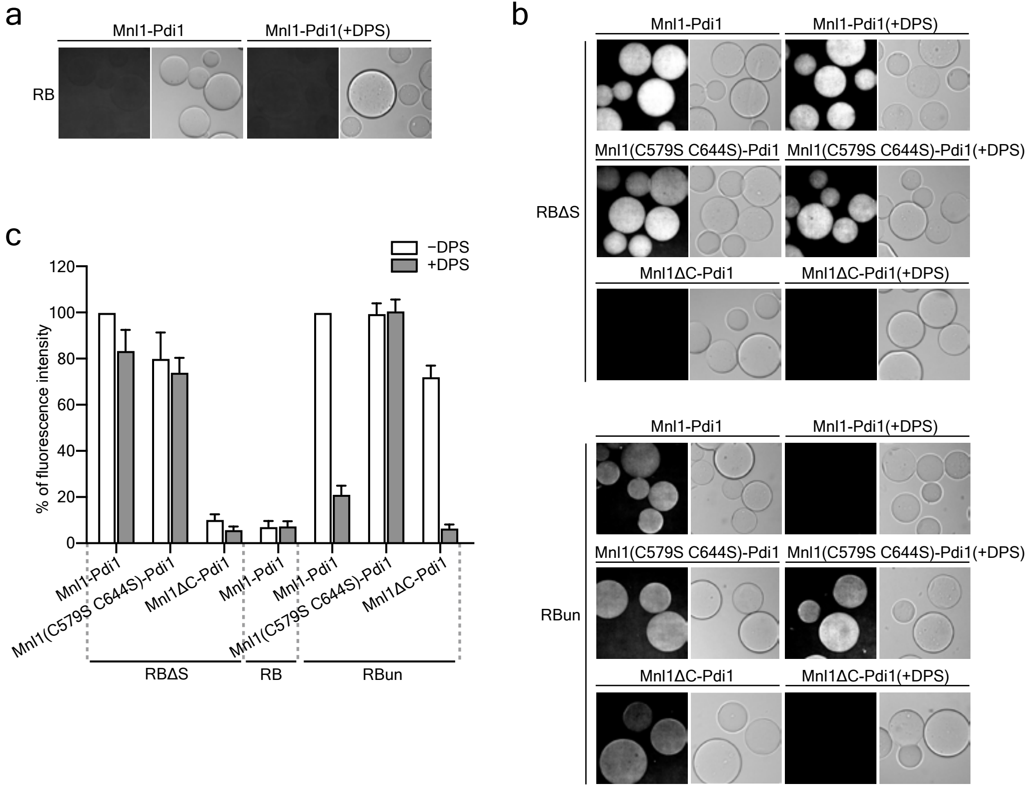

### ED figure 7

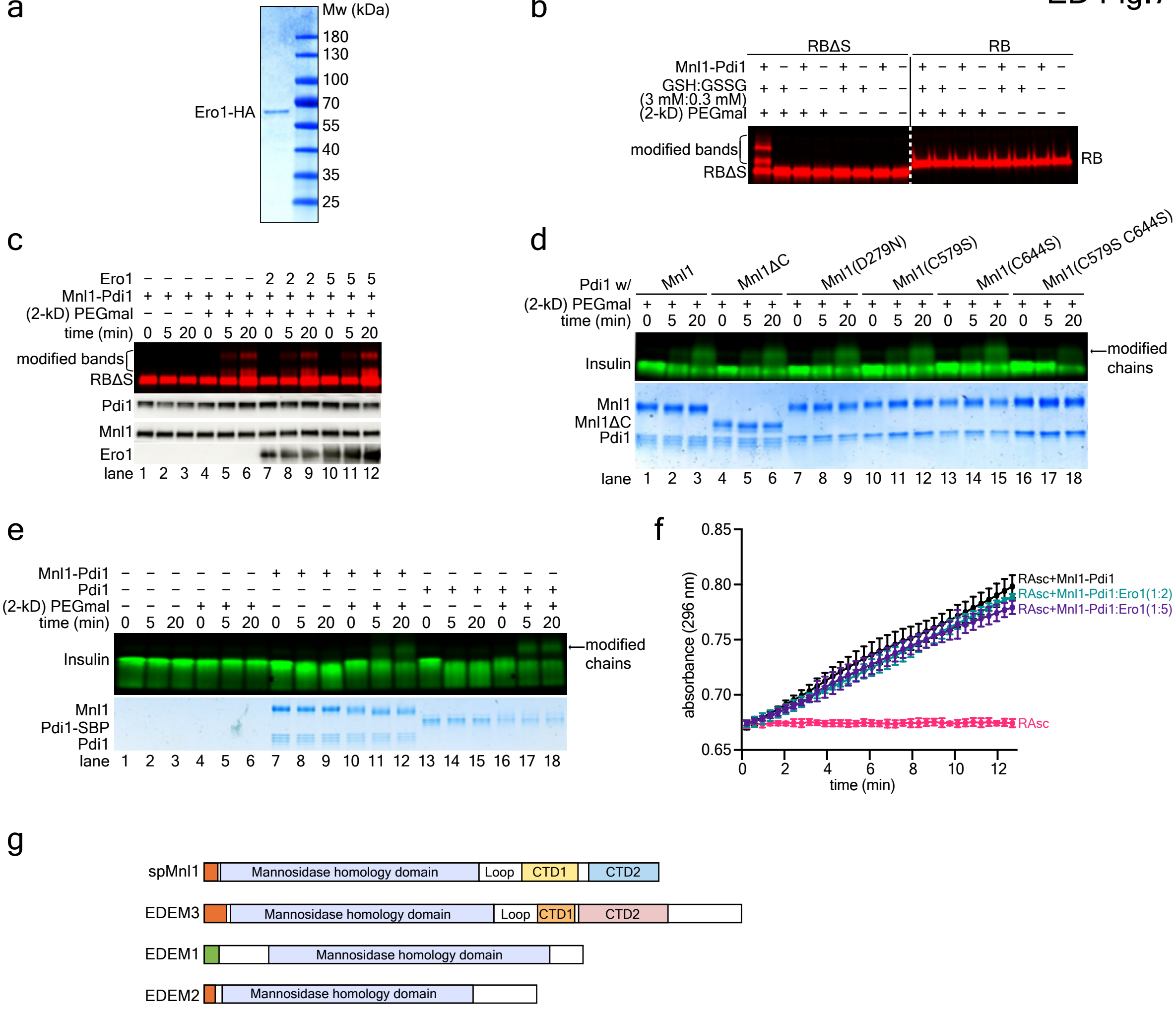

### Supplemental Figure 1

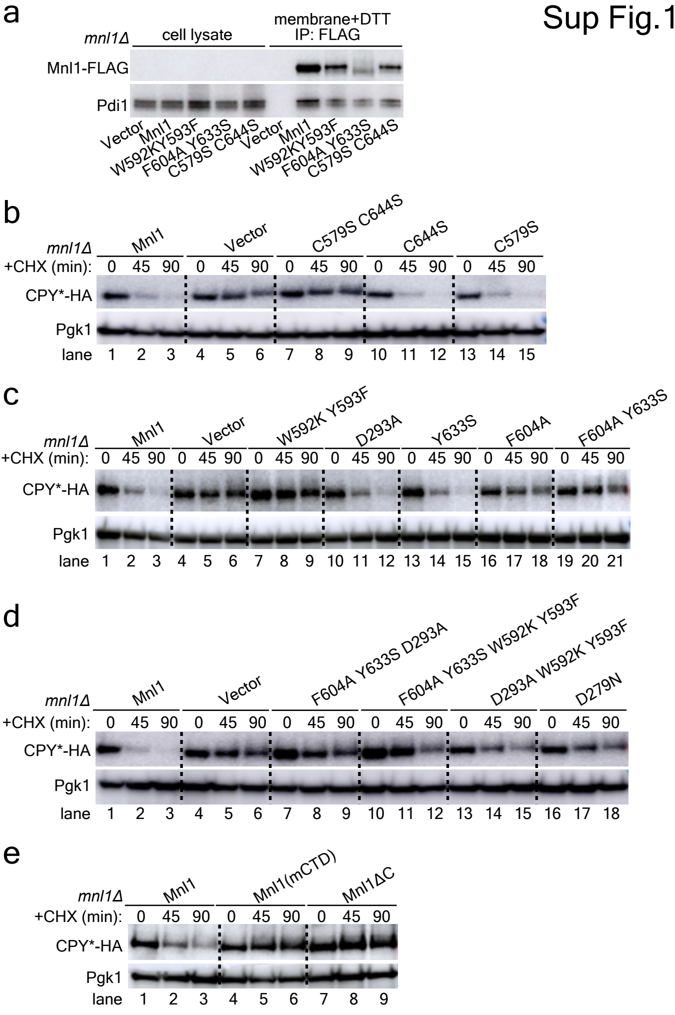

### Supplemental Figure 2

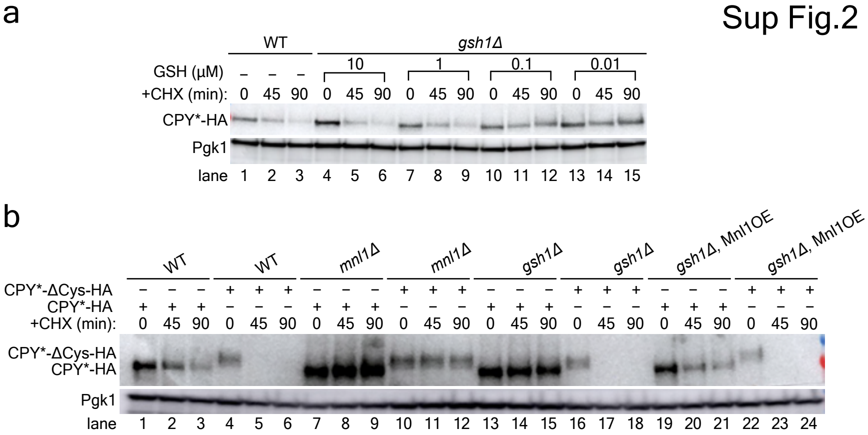
